# miRISC inhibition causes mitotic defects and synergizes with genotoxic agents in cancers

**DOI:** 10.64898/2025.12.12.693984

**Authors:** Minsi Zhang, Ziqi Jiao, Ylenia Cendon Florez, Ivana Lessel, Xiaoyi Li, Melissa A Yao, Moritz Weigl, Mateusz Ozimek, Turgut Dogruluk, Kevin Chen, Keith Conrad Fernandez, Vinagolu K. Rajasekhar, Olesja Popow, Joao A. Paulo, Davide Pradella, Tanmay Mishra, Chiara Mastroleo, Juliana I. Delgado, Francesco Enrico D’Amico, Vincenzo Cavalieri, Carol D. Morris, Meng-Fu Bryan Tsou, Kevin M. Haigis, Jayanta Chaudhuri, Robert Benezra, Joana A. Vidigal, Davor Lessel, Andrea Ventura, Gaspare La Rocca

## Abstract

Although individual microRNAs (miRNAs) can have tumorigenic or tumor-suppressive properties, their overall role in cancer remains controversial. Here, we show that cancer tissues and cell lines are characterized by preferential accumulation of the high-molecular-weight miRNA-induced silencing complex (HMWR), the functionally active form of the effector complex responsible for miRNA-mediated gene repression. Experimentally induced disassembly of the HMWR impairs the growth of human tumor xenografts and of autochthonous tumors in mouse models of human cancer *in vivo*. Furthermore, disassembly of the HMWR increases chromosome mis-segregation, which synergized with genotoxic agents to potentiate cancer cell vulnerability and improve therapeutic response. These findings suggest pharmacologic inhibition of HMWR as a novel anti-cancer strategy.

---

Metazoan microRNAs (miRNAs) are short (*∼*22nt-long) single-stranded RNAs that repress mRNAs by inducing their decay via partial base complementarity^1-3^. The human genome encodes hundreds of miRNAs^4,5^, often displaying highly tissue-specific expression patterns^6^. Each miRNA can bind to and regulate multiple mRNAs. Conversely, individual mRNAs often contain binding sites for several distinct miRNAs^6^. This complexity makes miRNA-mediated gene repression a versatile layer of post-transcriptional gene regulation^7^.

The mature form of miRNAs results from the processing of kilobases-long transcripts by the coordinated action of the miRNA biogenesis factors, which for most miRNAs includes the two endonucleases DICER and DROSHA, and accessory factors such as DGCR8, TRBP1 and XPO5, among others^8^. Once matured, miRNAs are loaded into the miRNA-induced silencing complex (miRISC), with Argonaute proteins (AGO1-4), and TNRC6 proteins (TNRC6A-C) as its core components^9,10^. AGO directly binds the miRNA and promotes base pairing with cognate targets^11^, while TNRC6 binds to AGO and acts as a scaffold for the recruitment of decapping and deadenylase complexes on target mRNAs^12-22^. This process results in mRNA decay, impaired translation and subsequent decreased protein output^2^.

Dysregulated miRNA function has been observed in several pathological conditions^23^, including cancer^24-27^. While individual miRNAs and miRNA families can have oncogenic or tumor suppressive properties, often in a context-dependent manner^28-33^, the consequences of global loss of miRNA function on cancer initiation and progression are unclear.

Efforts to address this question by measuring miRNA maturation and abundance in cancers have provided contradictory results. Several studies have described global down-regulation of miRNAs along with lower levels and impaired activity of their biogenesis factors in human cancers^34-52^. Intriguingly, opposite effects have also been observed^53-58^. The interpretation of these studies is further complicated by the observation that heterozygous deletion of Dicer is pro-tumorigenic, while its homozygous loss is tumor suppressive in mice^59-61^.

These conflicting observations may be explained by the fact that miRNA abundance and the activity of miRNA biogenesis factors are not reliable indicators of global miRNA function. Three lines of evidence support this notion. First, loss-of-function of genes involved in miRNA processing does not necessarily result in equal down-regulation of all miRNAs^40-42,46,62-64^ because some miRNAs bypass the canonical biogenesis pathway^65-71^ and mature miRNAs tend to be extremely stable^72,73^. Consequently, disruption of the miRNA biogenesis pathway is more likely to result in the enrichment of a subgroup of miRNAs within the miRNA pool, rather than in loss of global miRNA function^40-42,62,63^. Second, mounting evidence indicates that the miRISC is a dynamically regulated complex whose activity ultimately depends on intracellular and extracellular cues that regulate its assembly and target engagement^73-78^. Thus, elevated miRNA levels may not influence gene expression in the absence of active miRISC. Third, the interpretation of studies involving inhibition or deletion of genes controlling miRNA biogenesis is challenging because these genes additionally exert miRNA-independent functions^67,79-93^.

To overcome these limitations, we recently developed a strategy to inactivate the miRISC acutely and reversibly in cellular and murine models without affecting miRNA biogenesis or abundance^94^. Here we apply this approach to examine the consequences of chronic and acute loss of miRNA function on tumor initiation, progression, and maintenance in multiple cancer cell lines and mouse models of human cancers.

## Results

### miRNA function is controlled by miRISC assembly

We have previously shown that AGO proteins can be found as part of either a low molecular weight miRISC (LMWR) or a high molecular weight miRISC (HMWR), in a cell and tissue-dependent manner^73,75^. We have proposed that the HMWR represents the functional miRISC engaged in target repression, while the LMWR consists of AGO proteins bound to miRNAs, but temporarily disengaged from their target mRNAs^73,75^(Figure 1A, top panel). A key prediction of this model is that removal of an abundantly expressed miRNA should affects gene expression only if the HMWR is present. To formally test this hypothesis, we examined the consequences of targeted deletion of miR-92a^95^ in splenic B cells, a cell type in which a rapid and nearly complete transition of AGO proteins from LMWR to HMWR is observed upon mitogenic stimulation (Figure 1B, top panels). In agreement with our hypothesis, loss of miR-92a had no effect on the expression of miR-92a targets in resting B cell, where AGO proteins are predominantly in LMWR (Figure 1B, bottom left panel, Figure S1A, left panel and Dataset 1), while it resulted in their marked up-regulation in activated B cells, when the majority of AGO is part of the HMWR (Figure 1B, bottom right panel, Figure S1A, right panel and Dataset 1). Importantly, expression levels of miR-92a and other members of the miR-92 seed family did not change between the two conditions^90^ (Figure S1B). These results are consistent with the notion that the presence of HMWR indicates a functional miRISC engaged in target repression through the miRNA pathway. Furthermore, they offer a mechanistic explanation for prior observations by the Bartel group that targeted deletion of miR-150 had little or no impact on the expression of its predicted targets in resting mature B cells, despite being one of the most highly expressed miRNAs in this cell type^2^.

**Figure 1.**
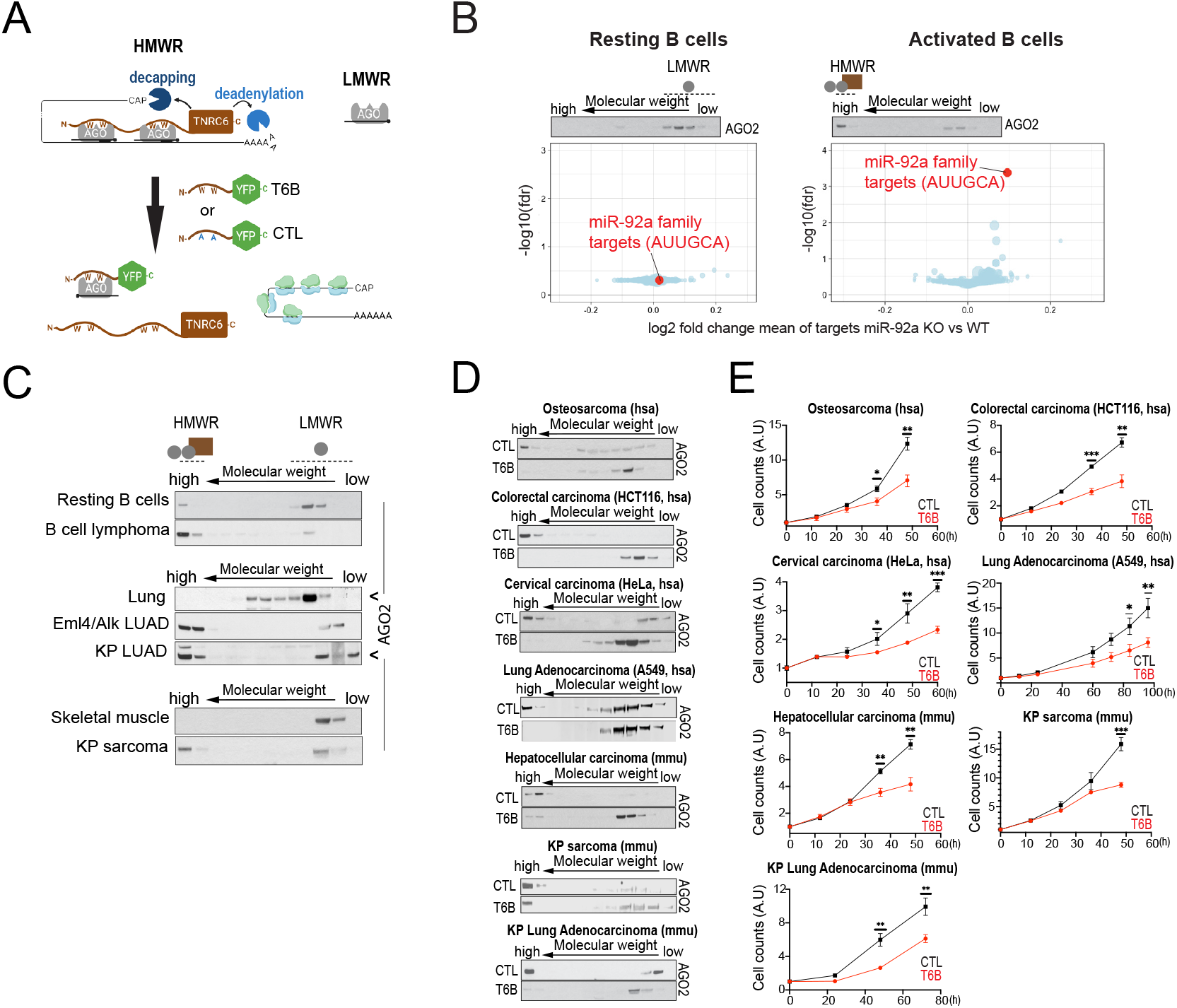
Inhibition of miRISC function is detrimental to cancer cell proliferation. **(A)** AGO proteins exist as part of a high molecular weight miRISC (HMWR, top left) engaged in target repression, or as part of a low molecular weight miRISC (LMWR) that lacks TNRC6 and is not associated to target mRNA (top right). Bottom, T6B peptide binds to AGO through tryptophan (W) residues and prevents its association with TNRC6, leading to de-repression of target mRNAs. A control (CTL) peptide has been designed to carry alanine residues (A) in place of the key tryptophan residues responsible for binding to TNRC6. **(B)** Top panels: Western blot showing the size exclusion chromatography (SEC) profile of AGO2 complexes in resting and activated B cells. AGO2 was used as a surrogate for T6B-mediated effects on all four AGO paralogues. Bottom panels: relative volcano plots of changes in expression of predicted miRNA targets for each conserved miRNA family upon miR-92a deletion. The indicated AUUGCA “seed” sequence characterizes miR-92a family members and confers target specificity. Note that miR-92a targets (red circles) are preferentially upregulated compared to the targets of other miRNA families only in activated B cells. For both panels, miRNA targets are grouped by circles, each representing a conserved miRNA family. The size of each circle is proportional to the number of predicted targets within the family. Two-sided Student’s t-test is used to compare matched WT and miR-92a KO groups. **(C)** Western blots showing SEC elution profiles of AGO2 complexes in normal tissues and in their malignant counterparts. **(D)** Western blots showing SEC elution profiles of AGO2 complexes of multiple cancer cell lines expressing either T6B or the control (CTL) peptide. **(E)** Growth curves of cancer cell lines as in (D).

Inhibition of miRISC impairs cancer cell proliferation *in vitro* The formation of the HMWR is dependent on TNRC6 expression levels, which are in turn induced by mitogenic signals^73,75,90,96^. Based on this observation, we reasoned that the assembly of the HMWR and consequent induction of miRNA function may be required during tumor development, a hallmark of which is sustained proliferation^97^. To investigate the dependency of tumors on global miRNA function, we employed size exclusion chromatography^75^ to compare the molecular size of the miRISC in a panel of primary mouse cancers and in their corresponding normal tissues. Eμ-Myc-driven B cell lymphomas^98^, lung adenocarcinomas (LUAD) driven by either Eml4-Alk^99^ or KRas^G12D^ ; p53^null^ (KP)^100^, and KP-driven sarcomas^101^, all showed striking accumulation of HMWR compared to their corresponding normal tissue in which the majority of AGO eluted in lower molecular weight fractions (Figure 1C). These findings prompted us to test the consequence of impaired miRISC on the proliferative potential of a variety of cancer cell lines, which invariably show AGO largely as part of the HMWR^75^. To impair miRISC function in these cell lines we expressed a transgene coding for T6B, a fragment of the AGO-binding domain of human TNRC6B that can disrupt the interaction between AGO and TNRC6 (Figure 1A, bottom panel)^78,94,102-109^. We also employed a control peptide (CTL) in which key tryptophan residues responsible for binding to AGO were substituted with alanine, thus rendering it inactive^94^. Both T6B and CTL cDNAs were fused in frame to HA and FLAG tags at the N terminus, and to either YFP or BFP at the C terminus (Table 1) and transduced into multiple cancer cell lines under the control of a doxycycline inducible promoter.

**Table 1.**
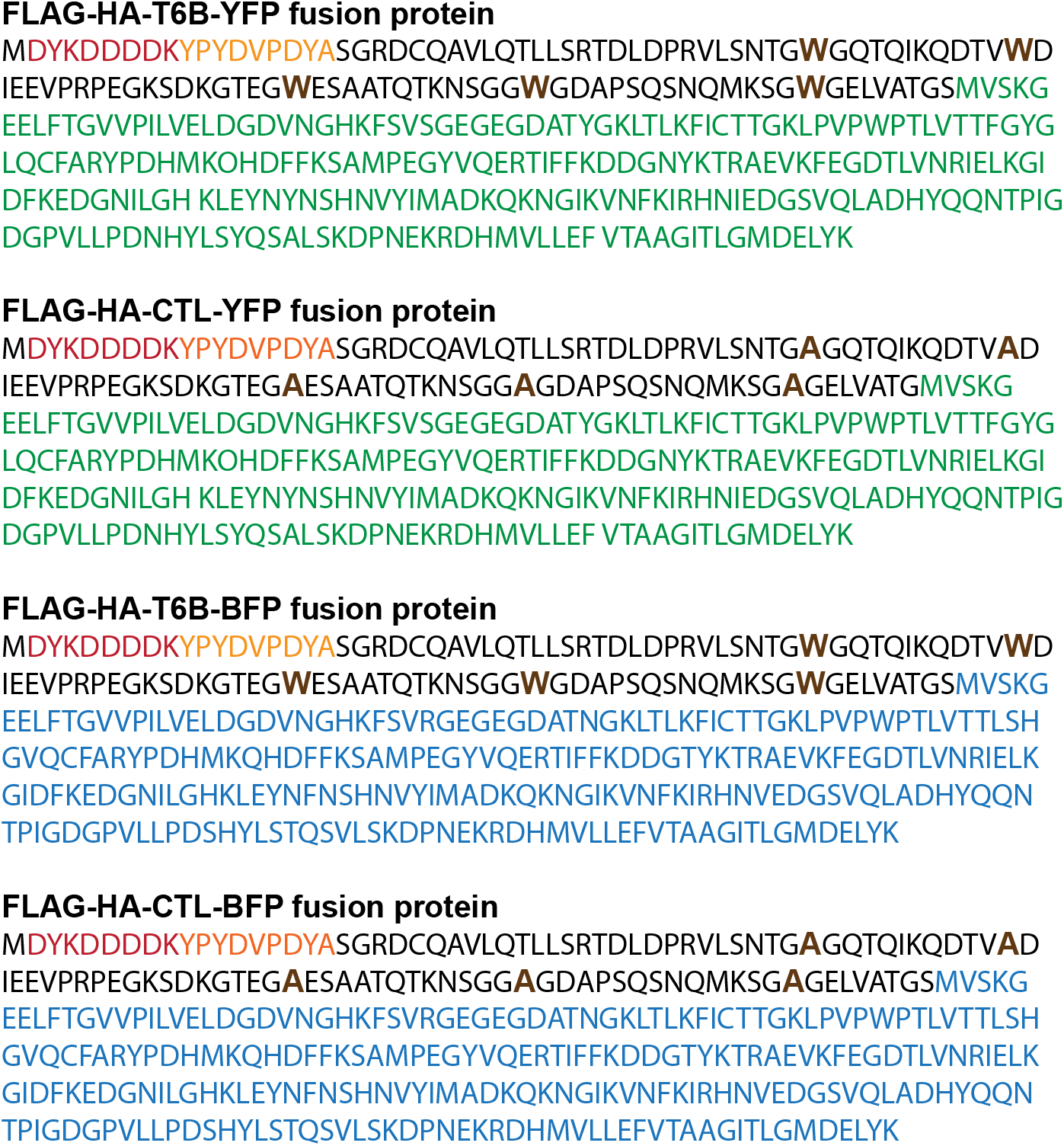
Amino acid sequence of the T6B and Control fusion proteins. Both fusion protein versions have HA and FLAG tags at the N termini and are fused to either the yellow fluorescent protein (YFP) or to the blue fluorescent protein (BFP) at the C-termini. In CTL all tryptophan residues (brown) are mutated to alanine to prevent interaction with AGO proteins. Red: FLAG-tag; orange: HA-tag; black: T6B; green: YFP. blue: BFP.

As expected, T6B expression induced dissociation of the majority of AGO from the HMWR (Figure 1D) in all tested cancer cell lines. Strikingly, it also markedly impaired their proliferation, while the CTL peptide failed to do so (Figures 1E and S1C). Importantly, T6B expression in a triple knock out (TKO) cell line for AGO1, 2 and 3, the most abundant AGO paralogues in vertebrates, did not affect cell proliferation (Figure S1D), indicating that the observed T6B effects are due to on-target impairment of AGO functions.

### Inhibition of miRISC impairs the growth of xenograft and autochthonous tumors *in vivo*

To extend these findings to a more physiological context, we assessed the consequences of impaired miRISC function on tumor growth in tumor xenograft. We subcutaneously injected A549 cells expressing either T6B-YFP or CTL-YFP transgenes in nude mice, maintained the mice on doxycycline chow, and monitored tumor burden over time (Figure S2A). T6B significantly reduced the growth of A549-derived tumors *in vivo* (Figures 2A and 2B). Interestingly, in all tumors that eventually developed in mice injected with T6B-YFP-expressing cells, expression of the T6B transgene was invariably lost while it was retained in tumors derived from cells expressing the CTL peptide (Figures 2C and 2D), indicating strong negative selection against T6B expression. We confirmed this result by size exclusion chromatography, which showed that decreased expression of T6B was accompanied by reconstitution of the HMWR (Figure 2E). We observed a similar selection against T6B but not against CTL expression in xenografts from HCT116 (Figures S2B and S2C) and HeLa cells (Figure S2D). These results demonstrate that inhibition of miRISC reduces miRNA function and is detrimental for the proliferation of cancer cell lines *in vivo* in a cell-autonomous fashion. We next investigated the consequences of inhibition of miRISC on the initiation and progression of autochthonous tumors in an immunocompetent context using genetically engineered mouse models of human cancers. To control T6B expression *in vivo*, we employed a genetically engineered mouse strain (*R26*^*rtTA*^ ; *Col1a1*^*T6B*^, hereafter R26^T6B^)^94^ in which a doxycycline-inducible T6B-YFP transgene is knocked into the *Col1A1* locus, and in which the rtTA transgene is constitutively expressed from the *Rosa26* (R26) locus (Figure S3, top panel). In this strain, doxycycline-induced T6B expression is well tolerated under homeostatic conditions^94^, thus enabling long-term studies on tumor development.

**Figure 2.**
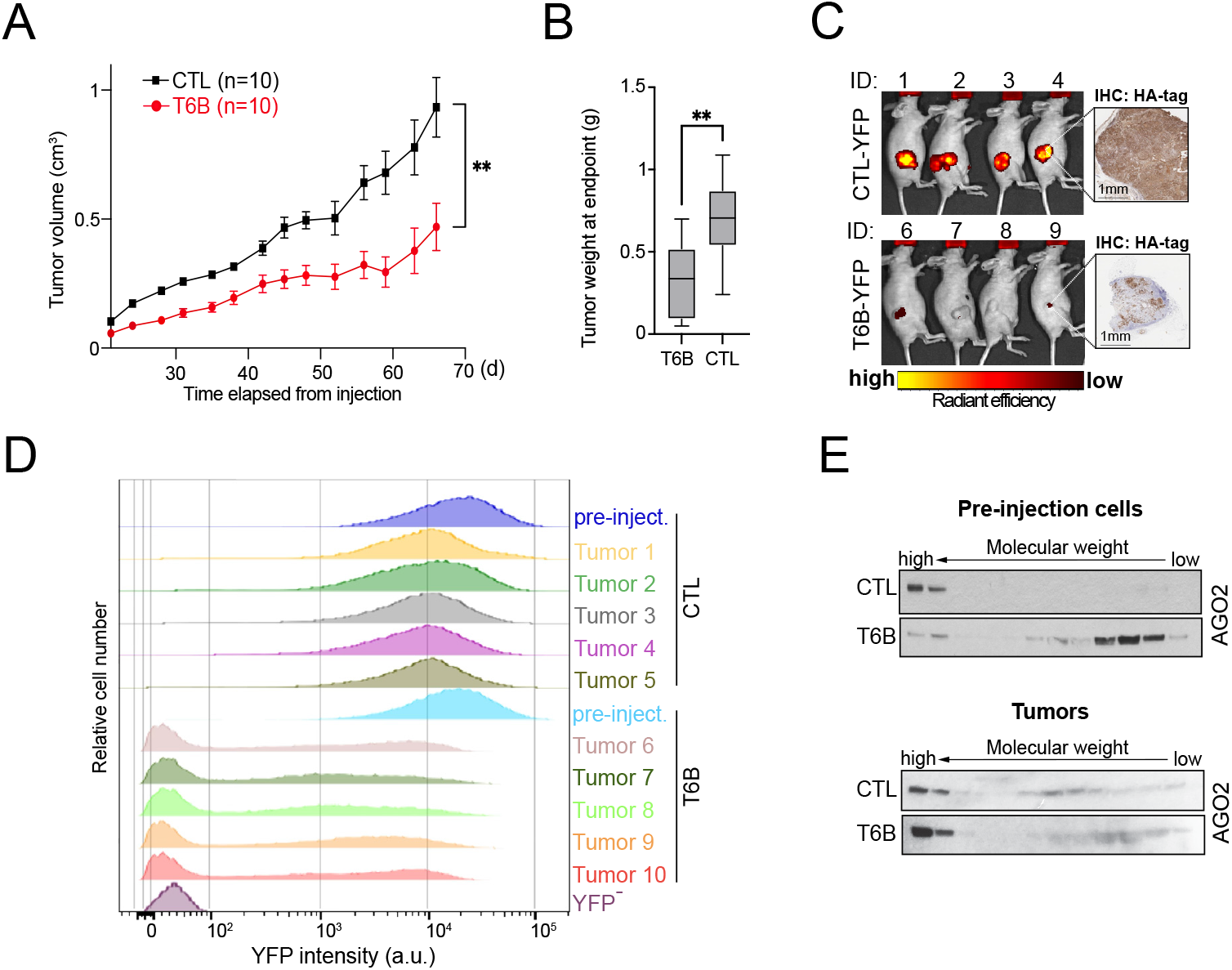
Inhibition of miRISC function impairs the growth of xenografts. **(A)** Growth curves of tumors derived from A549 cells expressing either T6B or the CTL peptide (n = 10 for each group). **(B)** Weight at endpoint of A549 tumors represented in (A). **(C)** Left, scans taken by *in vivo* imaging showing YFP fluorescence intensity in T6B- or CTL-expressing tumors derived from A549 cells at endpoint. Right, immunohistochemistry (IHC) staining using an antibody against the HA-tag showing the expression levels of T6B and the CTL peptide in sections of formalin-fixed/paraffin-embedded tumors. **(D)** Histogram plots showing distribution of YFP intensities, assessed by flow cytometry, in pre-injected T6B- and CTL peptide-expressing A549 cells and in cell suspension of right flank tumors at endpoint. Parental A549 cells (YFP-) served as negative control for both plots. **(E)** Western blots showing SEC profiles of AGO2 complexes in pre-injected A549 cells expressing either CTL or T6B peptide (top panels) and in cells derived from their tumors (bottom panels) at endpoint. Error bars indicate mean ±SEM in A, min to max range in B. ** p<0.01. Two-sided Student’s t-test is used to compare matched T6B and CTL groups. IHC: immunohistochemistry.

We induced the formation of LUAD driven by the *Eml4-Alk* gene fusion in R26^T6B^ using a CRISPR/Cas9-based strategy we previously developed^99^ that consists in intratracheal delivery of recombinant adenoviral vectors expressing both Cas9 and two guide RNAs designed to induce the 11 Mbp intrachromosomal inversion resulting in the *Eml4-Alk* rearrangement (Figure 3A, upper panel). As controls we used mice expressing rtTA but not the T6B-YFP transgene (*R26*^*rtTA*^; *Col1a1*^*wt*^, hereafter R26^rtTA^, Figure S3, bottom panel)^94^. As previously reported^99^, multiple bilateral lung tumors developed in both cohorts by two months post-injection, at which point we placed the mice on doxycycline-containing chow to induce T6B expression and disrupt the HMWR (Figure 3A and 3B).

**Figure 3.**
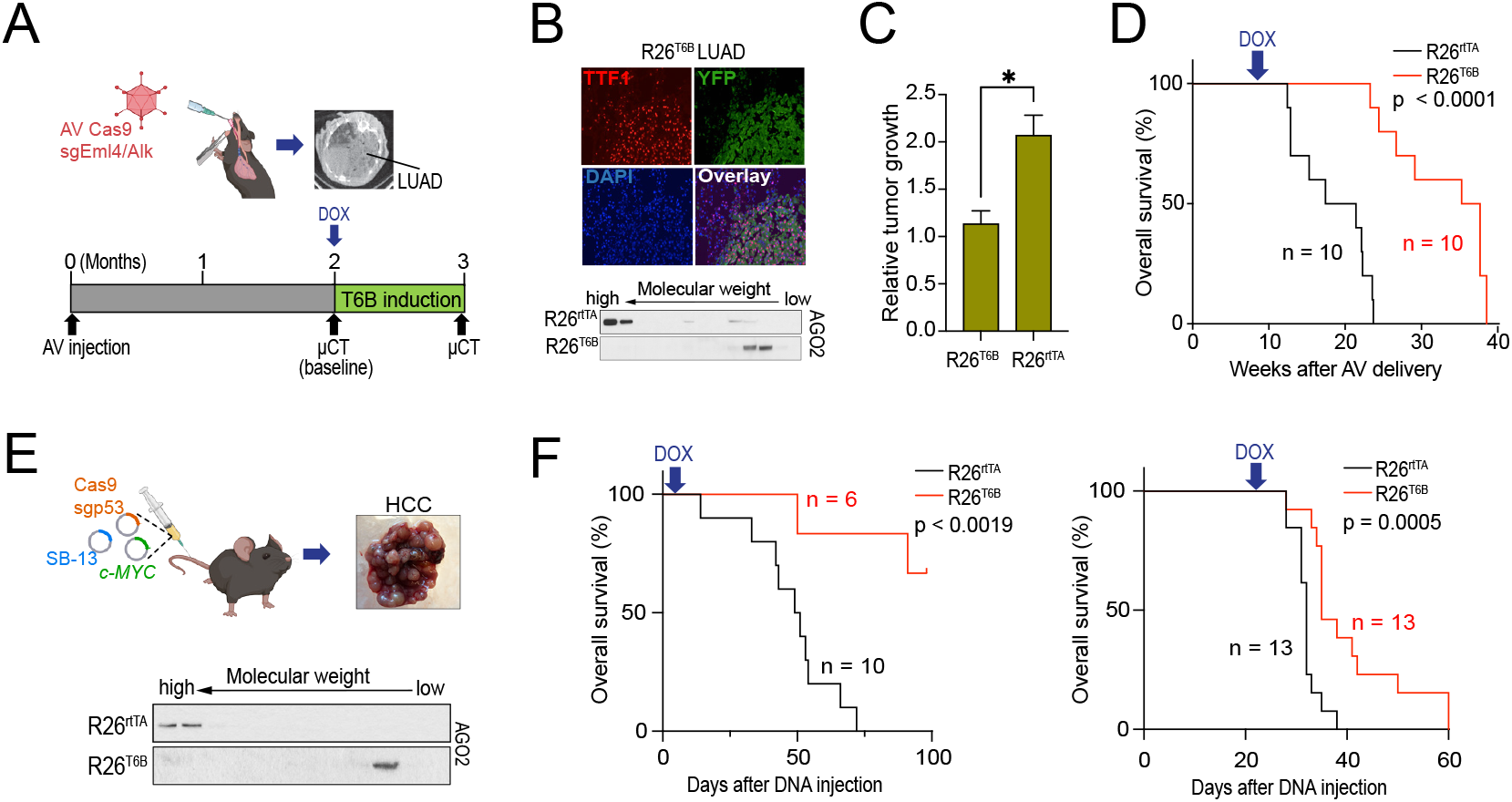
Inhibition of miRISC function reduces tumor growth and extends survival in autochthonous LUAD and HCC mouse models. **(A)** Top, representations of the procedure used to induce LUADs in in *R26*^*rtTA*^; *Col1a1*^*T6B*^ (R26^T6B^) and *R26*^*rtTA*^; *Col1a1*^*wr*^ (R26^rtTA^) mice by Cas9-medaited *Eml4/Alk* genomic rearrangement. Bottom, experiment timeline for tumor induction and detection. Lung images shown on top right of the figure were generated by μCT scan. **(B)** Top, immunofluorescence images showing the expression of T6B fusion protein and of the thyroid transcription factor 1 (TTF1), a diagnostic marker for lung cancer, in sections of fresh-frozen *Eml4-Alk* LUADs generated in R26^T6B^ mice. T6B expression was detected by using an antibody against YFP. Note the colocalization of T6B and TTF1 signals. Bottom, Western blots showing SEC profiles of AGO complexes in *Eml4-Alk* LUADs generated either in R26^T6B^ or R26^rtTA^ mice. **(C)** Box and whisker plots of relative volume increase of LUADs 1 month after doxycycline administration. Tumor volume measurements were conducted on μCT scans (n = 6 for CTL, n = 3 for T6B) and were normalized to tumor density at baseline, as schematized in (A). **(D)** Survival curves for cohorts of R26^T6B^ and R26^rtTA^ mice harboring *Eml4-Alk* LUADs. **(E)** Top, schematics of the procedure adopted to generate hepatocellular carcinomas (HCC) in R26^T6B^ and R26^rtTA^ mice and representative image of the resulting hepatic tumors. Bottom, Western blots showing SEC elution profiles of AGO2 complexes in CTL and T6B tumors. **(F)** Survival curves for cohorts of R26^T6B^ and R26^rtTA^ mice harboring *Myc/p53null* HCC tumors. Doxycycline administration started either on day 4 (left panel), or on day 25 (right panel) post tumor initiation. SB-13: sleeping beauty transposon. sgp53: single guide against murine *Trp53*. Log rank-test is used to compare matched T6B and CTL groups in the respective Kaplan Meier survival curves.

Thirty days after initiation of doxycycline administration, tumor burden in control tumors had doubled as compared to baseline, while T6B-expressing tumors displayed negligible changes (Figure 3C). Consistent with this observation, doxycycline administration resulted in an increased overall survival of R26^T6B^ animals compared to controls (Figure 3D).

To validate these observations in a different tumor type, we induced hepatocellular carcinomas (HCC) in R26^T6B^ and control mice, employing a previously described method based on the somatic delivery by hydrodynamic tail vain injection (HDTVi) of plasmids designed to inactivate *Trp53* by CRISPR-Cas9 and integrate a constitutively-expressed *c-Myc* oncogene by sleeping beauty transposase (SB)^110,111^. These two genetic events are sufficient to drive the formation of multiple HCCs within few weeks^110^ (Figure 3E, upper panel). Importantly, in these tumors nearly all AGO2 shifted from HMWR to LMWR upon T6B expression (Figure 3E, lower panel).

To explore the consequences of miRISC disruption during the early vs late stages of tumorigenesis, we induced T6B expression at different time points after HDTVi. In a first cohort, doxycycline was administered early on at 4 days post-injection, while in the second cohort it was administered 25 days post-injection, when tumors are already detectable. We monitored animals of both cohorts for the development of tumors and overall survival. In both experimental settings, the R26^T6B^ animals showed an extended overall survival compared to the control animals (Figure 3F). Interestingly, in the cohort where doxycycline administration started early after HDTVi, while all animals of the control group had succumbed within 75 days, 4 out of the 6 R26^T6B^ animals were still alive and tumor free at 100 days after HDTVi (Figure 3F, left panel).

Collectively, these results indicate that inhibition of miRISC results in impaired growth of autochthonous tumors in mice, both at early and late stages of tumorigenesis.

### Inhibition of miRISC results in aberrant mitosis regulation and promotes genomic instability

To gain mechanistic insights into how miRISC inhibition affects cancers, we analyzed gene expression by RNAseq in T6B- and CTL-expressing HCT116, KP sarcoma and KP LUAD cell lines. As expected, T6B expression resulted in preferential upregulation of predicted miRNA targets in all three cell lines (Figures 4A, Figure S4A and Dataset 2), confirming inhibition of miRNA-mediated gene repression in these cells. Cross-comparison of mRNA expression levels identified a core of 208 upregulated (Figure 4B, top panel) and 130 downregulated genes (Figure 4B, bottom panel and Dataset 3) shared by all tested cancer cell lines. In agreement with the role of T6B in derepressing miRNA targets globally, miRNA-targets enrichment analysis^112^ identified a significant enrichment of miRNA-target interactions within the common upregulated genes, but not in the downregulated genes (Figure S5 and Dataset 4). Over-representation analysis (ORA) of the common upregulated genes showed significant enrichment for terms related to cell cycle and mitotic regulation such as “mitotic spindle” and “G2/M checkpoint” (Figure 4C, top panel), suggesting dysregulation of the mechanisms controlling these cellular processes. Conversely, ORA of the common downregulated genes identified “DNA repair” and “oxidative phosphorylation” as the most enriched terms (Figure 4C, bottom panel and Dataset 3).

**Figure 4.**
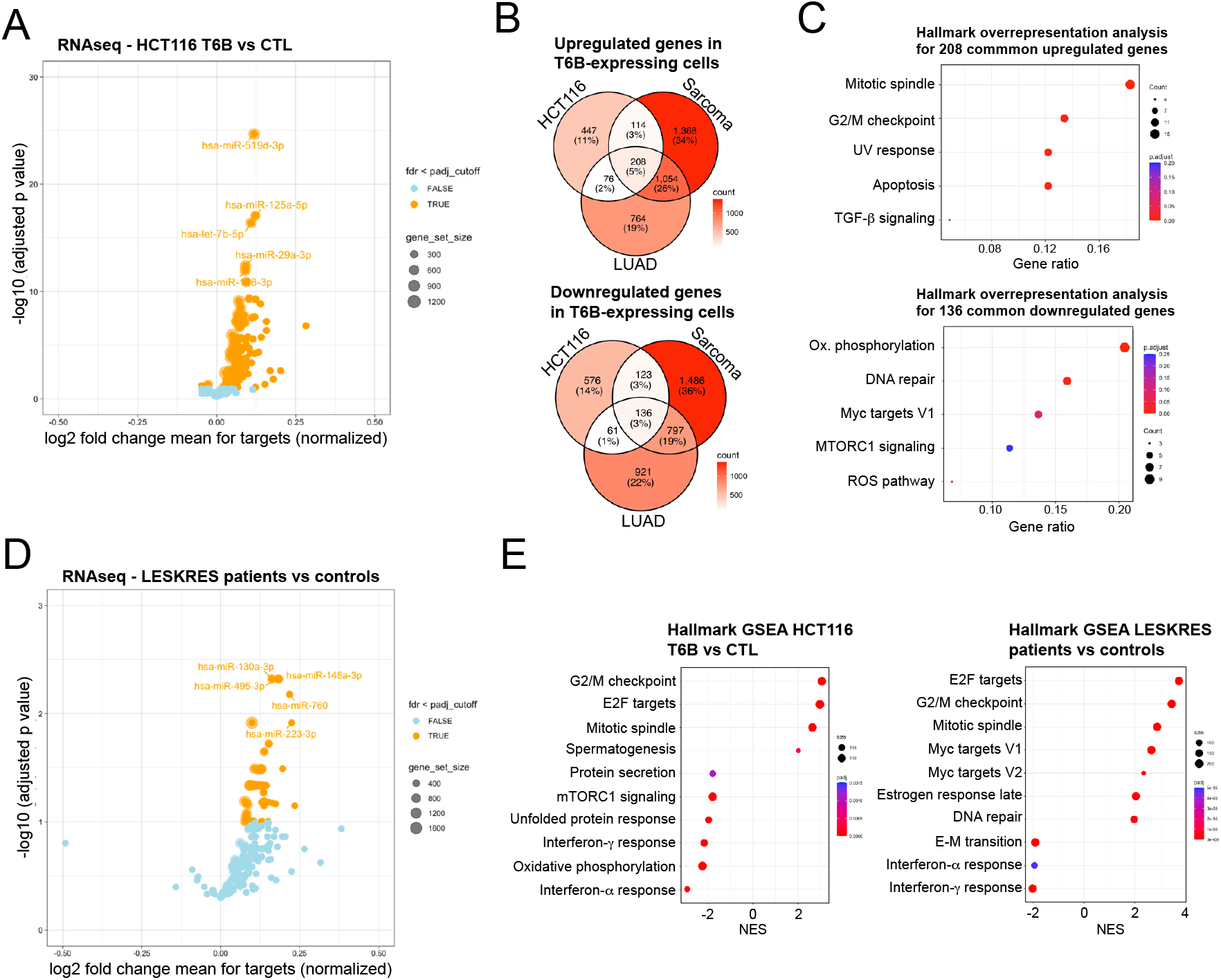
T6B expression recapitulates gene expression signatures caused by AGO2 mutations. **(A)** Volcano plots of mRNAs showing preferential global up-regulation of predicted miRNA targets for each conserved miRNA family in HCT116 cells expressing T6B compared to cells expressing the CTL peptide. miRNA targets are grouped by circles, each representing a conserved miRNA family. The size of each circle is proportional to the number of predicted target genes within a given miRNA family. **(B)** Venn diagram of genes that are commonly upregulated (left) and down-regulated (right) in all three sequenced T6B tumor cell lines. **(C)** Over-representation analysis (ORA) of commonly upregulated (left) and downregulated (right) genes in (B) using annotated Hallmark gene sets. Top 5 differentially enriched terms are shown for each gene list. **(D)** Same analysis as in (A) for LESKRES patient-derived fibroblasts compared to matched controls. **(E)** Top: Gene Set Enrichment Analysis (GSEA) of differentially expressed genes in HCT116 cells expressing T6B compared to CTL using annotated Hallmark gene sets. Top 10 differentially enriched terms are shown. Bottom: same GSEA analysis for LESKRES patient-derived fibroblasts compared to matched controls.

It is possible that the gene expression signatures and proliferative defects observed in cancer cells upon T6B expression result from off-target activities that are not related to its binding to AGO proteins. To test this possibility, we sought an independent experimental system that, like in T6B-expressing cells, could recapitulate global inhibition of miRNA function. We reasoned that, if T6B repressed miRNA function specifically, such a system would phenocopy the transcriptional and cellular defects seen in T6B-expressing cells. The Lessel-Kreienkamp syndrome (LESKRES)^113^, a rare developmental syndrome caused by heterozygous AGO2 mutations, provided such opportunity, as LESKRES mutations have been found to cause aberrant AGO2-target interaction, which conceivably leads to global impairment of miRNA function. Indeed, analysis of RNA-seq data of LESKRES patient-derived fibroblasts^113^ showed preferential de-repression of predicted miRNA targets (Figure 4D and Dataset 2). Strikingly, gene set enrichment analysis (GSEA) revealed a remarkable concordance of terms related to cell cycle and mitotic regulations (Figures 4E and Dataset 5). Moreover, the most significantly upregulated genes in T6B-expressing HCT116, KP Sarcoma and KP LUAD cells were also preferentially upregulated in LESKRES cells (Figure S4B and Dataset 6). Collectively, these observations are consistent with the hypothesis that T6B expression affects mitosis through impairing Argonaute function.

To further assess to which extent miRISC impairment affects mitotic control mechanisms, we investigated interphase cells for mitotic abnormalities, including the presence of bi- and multinuclear cells, as well as CREST-negative and CREST-positive micronuclei. CREST-positive micronuclei reflect the presence of centromeric sequences, thus indicating chromosome mis-segregation^114^. We focused on HCT116 cells for high-resolution analysis due to their genomic stability and near-diploid karyotype, which makes mitotic abnormalities more easily detected over background. T6B expression resulted in higher occurrence of micronuclei, bi- and multinucleated cells (Figures 5A and 5B), indicating greater genomic instability. Importantly, whereas the frequency of CREST-negative micronuclei was comparable between T6B- and CTL-expressing HCT116, a significant increase was observed for CREST-positive micronuclei (Figures 5A and 5B). Crucially, we observed a similar increase of mitotic abnormalities— including elevated CREST-positive micronuclei and binucleation—in AGO2-mutant fibroblasts derived from LESKRES patients (Figures S6A and S6B).

**Figure 5.**
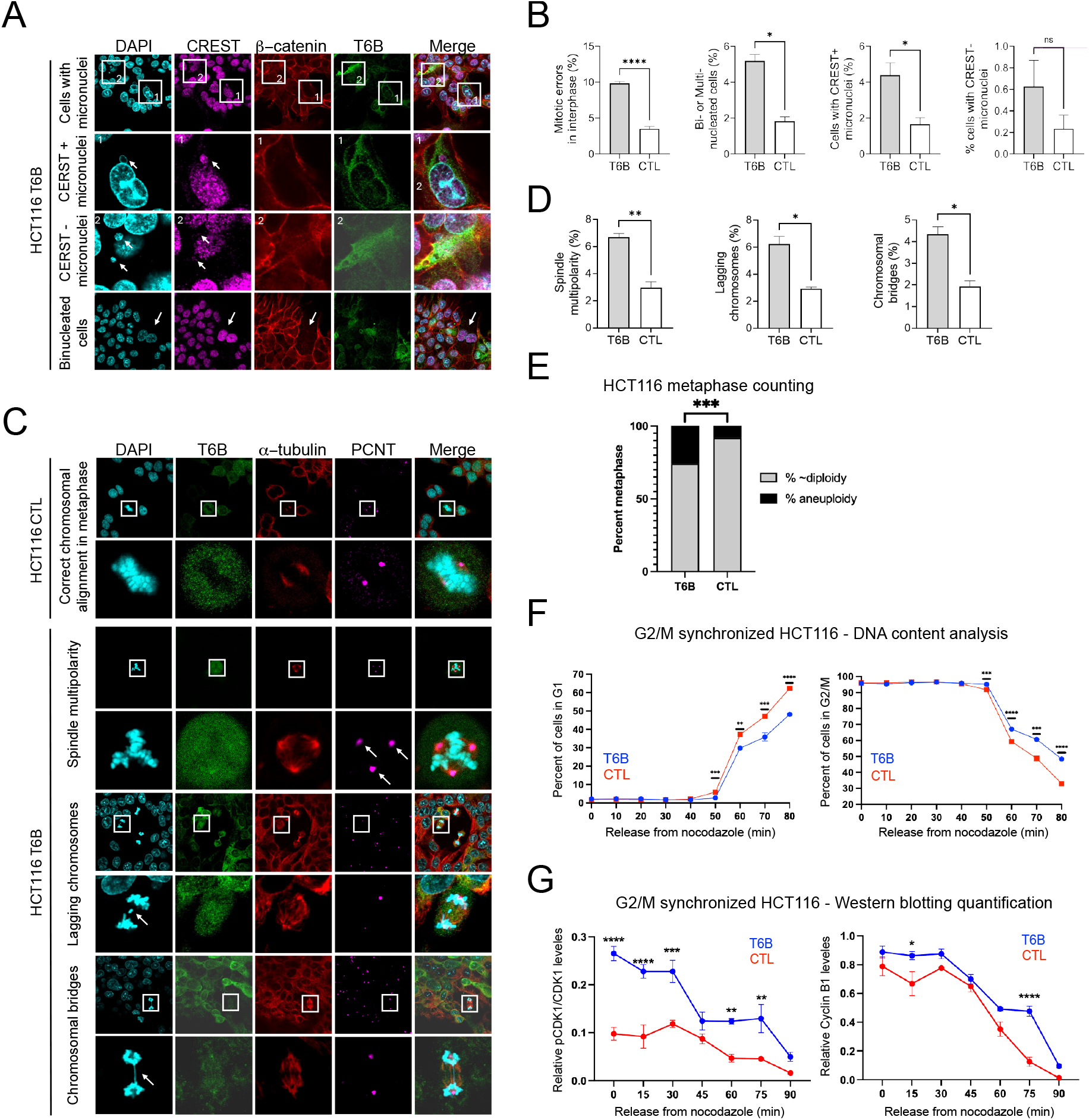
Inhibition of miRISC function results in defective mitosis in cancer cells and LESKRES patient-derived fibroblasts. **(A)** Phenotypic characterization of T6B-expressing HCT116 cells in interphase. Cells with multiple nuclei (bottom) and CREST+ micronuclei (1.) were observed and distinguished from cells with CREST-micronuclei (2.). **(B)** Quantification of specific cellular phenotype shown in (A) in at least 1500 cells per experiment. **(C)** Aberrant mitotic phenotype (multipolar spindle, lagging chromosomes and chromosomal bridges) of T6B- and CTL-expressing HCT116 cells were analyzed during mitosis. **(D)** Quantification of (C) with at least 100 mitotic cells analyzed per experiment. **(E)** Quantification of aneuploidy in T6B- or CTL-expressing HCT116 cells based on chromosomal spreads. **(F)** DNA content in 2N (left) and 4N (right) during mitosis analyzed using flow cytometry in G2/M synchronized T6B- or CTL-expressing HCT116 after nocodazole release. (G) Quantification of Western blots shown of Cyclin B1 (right) and Y15 phospho-CDK1 (left) during mitosis in G2/M synchronized T6B- or CTL-expressing HCT116 after nocodazole release. For each protein, tubulin-normalized quantifications are shown with representative blots on top. Data are represented as mean ± SEM. * P<0.05, ** p<0.01, ***p<0.001, ****p<0.0001. Two-tailed Student’s t-test is used to compare matched T6B and CTL groups in (B), (D), and (F). Fisher’s exact test was used to compare T6B and CTL metaphases in (C). Two-way ANOVA with Sidak’s correction is used to compare matched T6B and CTL timepoints in (G).

To gain additional insight into the origin of accumulation of mitotic defects, we analysed the frequency of multipolar spindles, lagging chromosomes and chromosomal bridges in HCT116 cells. We observed that T6B expressing cells displayed a significant increase in all these mitotic defects (Figures 5C, 5D), suggesting that miRISC disruption affects multiple aspects of mitosis^115-117^. Notably, T6B-expressing HCT116 cells and LESKRES patient-derived cells showed similar enrichment of Gene Ontology (GO) terms related to chromosome segregation and the kinetochore (Figure S7 and Dataset S7), highlighting the molecular and phenotypic convergence on chromosome mis-segregation in miRISC-impaired cells.

Because chromosomal segregation errors can promote aneuploidy^118^, we finally performed a chromosomal karyotyping analysis. As anticipated, we observed a significant increase in frequency of aneuploidy in HCT116 expressing T6B as compared to control (Figures 5E and S8).

Together, our results indicate that compromised miRISC function disrupts mitotic fidelity, predisposing cells to chromosomal mis-segregation and instability.

### Aberrant G2/M transition and mitosis in cancer cells under miRISC inhibition results in impaired cellular proliferation

Abnormal mitotic division and genomic instability can activate cell cycle surveillance and checkpoint pathways, in turn reducing the ability of cells to proliferate^119-121^. Indeed, we observed that T6B expression in HCT116 cells synchronized to G2/M boundary resulted in prolonged G2/M progression and delayed G1 re-entry (Figures 5F and S9A). These abnormalities are compatible with defects in mitotic timing arising from delayed mitotic entry or exit from mitosis, or a combination of both. To explore these possibilities, we measured mitotic progression by live cell imaging in T6B-expressing HCT116 cells and in hepatocellular carcinoma cells derived from autochthonous murine tumors (Figure 3E). Specifically, we measured the interval between rounding onset, which indicates entry into mitosis, and appearance of the cleavage furrow, which indicates completion of anaphase. These analyses showed that in both cell lines T6B expression led to significantly prolonged mitotic progression duration compared to control (Figure S9B). We also measured CDK1-phosphorylation (pY15-CDK1), a marker of G2/M checkpoint activation^122^ along with cyclin B1 degradation, a key molecular marker of metaphase to anaphase transition^123^. T6B expression in HCT116 cells synchronized to G2/M boundary resulted in enhanced and prolonged CDK1 (Tyr15)-phosphorylation (Figure 5G left and Figure S10), consistent with activation of the G2/M checkpoint and delayed mitotic entry. In addition, we observed a significant delay of cyclin B1 degradation in T6B expressing cells indicating dysregulated metaphase-to-anaphase progression (Figure 5G right and Figure S10). Together, these data suggest that miRISC inhibition triggers a G2/M checkpoint response, delays mitotic entry through sustained CDK1 inhibition, and impairs mitotic progression, likely contributing to the overall proliferation defect we observed in cancer cells upon miRISC inhibition.

### Inhibition of miRISC sensitizes cancer cells to genotoxic agents

Genotoxic agents—including radiation and chemotherapy— are mainstays of cancer treatment due to their ability to induce DNA damage and trigger mitotic catastrophe in proliferating cells^124^. Because miRISC-impaired cancer cells exhibit several mitotic abnormalities and elevated chromosomal instability, we hypothesized that these vulnerabilities could synergize with genotoxic agents.

To test this hypothesis, we assessed the response of a panel of T6B- and CTL-expressing cancer cells to a single dose of ionizing radiation. To avoid confounding effects caused by the antiproliferative activity of prolonged T6B expression, we pulsed the cells with doxycycline for only 24 hours prior to radiation to transiently induce the expression of the T6B or CTL peptide. Transient T6B expression significantly enhanced radiosensitivity across all tested cancer cells regardless of species or tissue of origin (Figure 6A, S11A and S11B) compared to cells expressing the CTL peptide.

**Figure 6.**
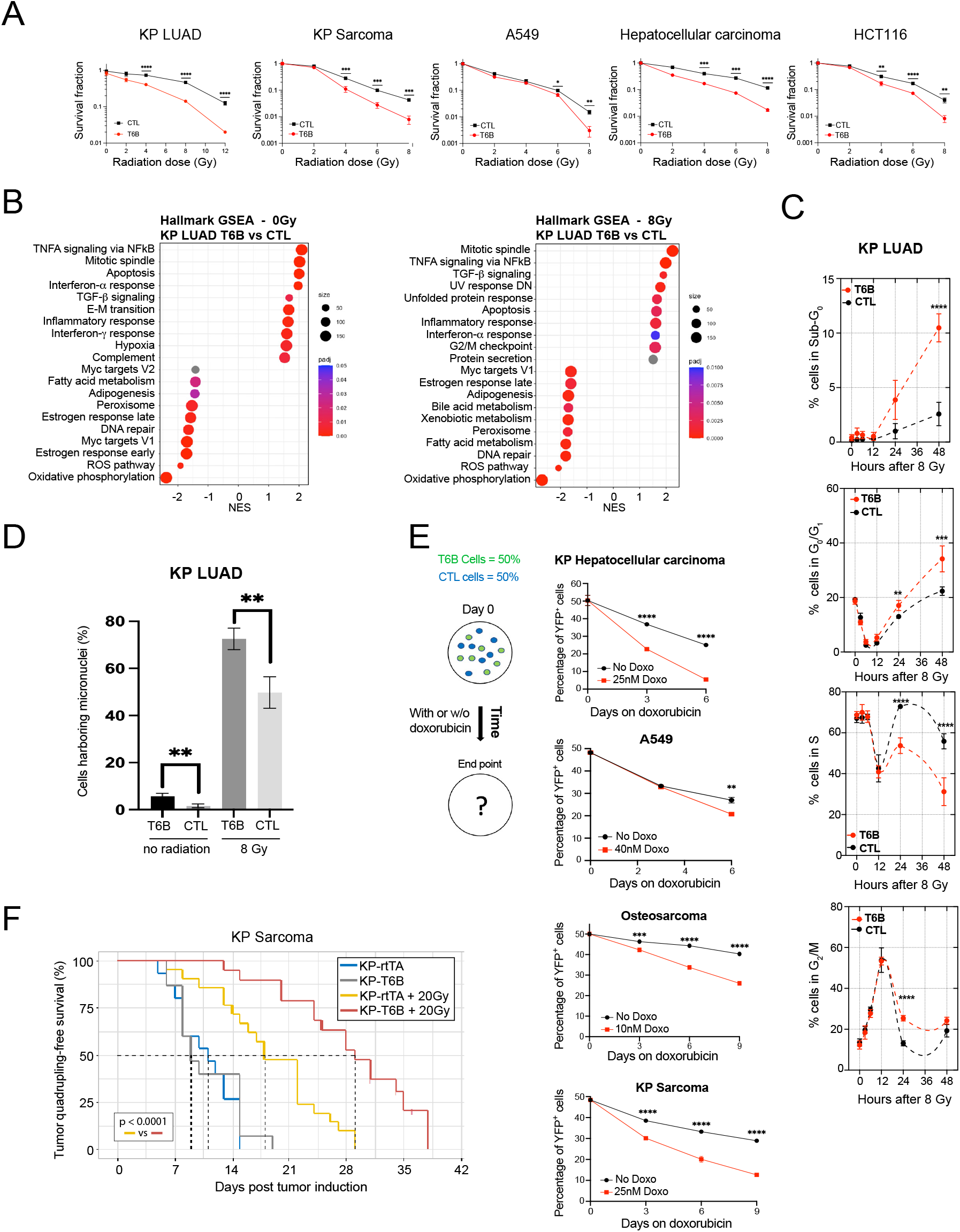
Inhibition of miRISC function enhances tumor sensitivity to genotoxic agents. **(A)** Colony formation assay showing response of T6B- and CTL-expressing cancer cells to a single dose of ionizing radiation at various intensities, as indicated in the x axis. **(B)** Gene Set Enrichment Analysis (GSEA) of differentially expressed genes in T6B vs. CTL KP LUAD cells receiving 0 (left) and 8gy radiation (right) using annotated hallmark gene sets. Top 10 differentially enriched terms in both directions are shown. **(C)** Distribution of percentages of KP LUAD cells in S, G2/M, sub-G0, and G1 phases over time after a single 8 Gy dose of radiation with a prior 24-hour pulsed expression of either T6B (black) or CTL peptide (red). **(D)** Percentage of micronuclei assessed by microscopy in T6B- and CTL peptide-expressing KP LUAD cells without radiation and after receiving a single dose of 8 Gy. **(E)** Top, schematic of competitive cell growth assay with or without doxorubicin treatment. Cells expressing the CTL peptide tagged by BFP are marked blue. Cells expressing T6B peptide tagged by YFP are marked green. Bottom, percentage of T6B (YFP+) cells remaining in culture over the course of experiment (n = 3 per timepoint). For each cancer type, results are representative of 3 independent experiments. **(F)** Tumor quadrupling-free survival curves for the indicated cohorts of KP-T6B and KP-rtTA mice bearing KP sarcomas, treated or not with a single 20 Gy dose of radiation. Error bars indicate mean ±SD. * P<0.05, ** p<0.01, ***p<0.001, ****p<0.0001. Two-sided Student’s t-test is used to compare matched T6B and control groups. Two-sided Student’s t-test is used to compare matched T6B and CTL in 6A, 6C, and 6E. Log-ranked test is used to compare matched to KP-rtTA and KP-T6B survival curves in 6F.

To explore the consequences of ionizing radiation on miRISC-impaired cancer cells, we focused on KP LUAD cells exposed to 8 Gy of ionizing radiation, a dose that severely reduced survival in T6B-expressing cells but was well-tolerated by control cells (Figure 6A and S11A). GSEA confirmed that T6B expression alone resulted in an enrichment of terms involved in mitotic regulation, which became even more significant upon radiation (Figure 6B and Dataset 8).

T6B-expressing cells displayed prolonged G2-M arrest in response to radiation compared to CTL-expressing cells. By 24 hours post-irradiation, T6B cells also displayed a marked increase in sub-G0 population—indicative of DNA fragmentation—which increased further at 48h (Figure 6C). This was accompanied by an increase in radiation-induced micronuclei formation compared to control cells (Figure 6D).

To extend our finding to a different source of genotoxic stress, we performed competitive cell growth assays using co-cultures of cancer cells expressing either T6B-YFP or CTL-BFP (Figure 6E) in the presence of doxorubicin, an intercalant agent that primarily causes DNA double-strand breaks and is widely employed in the treatment of diverse malignancies. Also in this context, genotoxic stress strongly synergized with T6B expression in impairing tumor cell growth (Figure 6E).

Encouraged by these results, we next tested the effects of genotoxic agents on the growth of T6B-expressing autochthonous *Kras-G12D; p53-null* (KP) soft tissue sarcomas, a mouse model of human cancer that has been used extensively to study radiotherapy and can tolerate high doses of radiation^125-127^. We generated a cohort of mice (KP-T6B) harboring the doxycycline-inducible T6B transgene in which CRE delivery leads to simultaneous activation of the *Kras-G12D* allele and deletion of the tumor suppressor *tp53*. We also generated a control cohort lacking the T6B transgene (KP-rtTA). We injected mice from both cohorts with CRE-encoding adenoviruses (AdCre) to induce the formation of sarcomas, which became detectable as early as 5 weeks post-injection, a time point at which we initiated doxycycline administration. Interestingly, although T6B induced miRISC disassembly in these sarcomas (Figure S11C), their growth was not significantly affected compared to controls (Figure 6F), even though cell lines derived from these tumors had impaired proliferative potential in culture (Figure 1E and S1C) following T6B expression (Figure S4A). This discrepancy between *in vitro* and *in vivo* observations may be due to insufficient expression of T6B *in vivo* to maintain robust and homogeneous miRISC inhibition, a possibility supported by the persistence of some HMWR in T6B tumors (Figure S11C). Nevertheless, progression free survival following radiation was significantly extended in the KP-T6B cohort compared to the KP-rtTA (Figure 6F) cohort.

These results indicate that inhibition of miRISC can broadly sensitize cancer cells to different DNA damage-inducing agents commonly employed as a component of standard of care for a variety of cancers, exemplified by enhanced radiosensitivity of sarcomas *in vivo*.

## Discussion

Here, we examined the role of the miRISC in tumor development. This study was motivated by the observation that various cancer cell lines and autochthonous tumors displayed a striking enrichment of HMWR compared to normal tissues, leading us to hypothesize that cancer cells heavily rely on global miRNA function. The ability of the T6B system to acutely disrupt the miRISC complex enabled us to formally test this hypothesis.

An essential role of miRISC in sustaining cell proliferation and tumor growth is supported by multiple and orthogonal experimental settings here employed: (i) 2-D cultures of multiple human and murine cancer cell lines revealed significant cytostatic effects due to T6B expression; (ii) Xenograft experiments showed that tumors derived from T6B-expressing cells exhibited markedly reduced growth, and those that did emerge, had lost T6B expression, indicating negative selection against miRISC inhibition; (iii) Several autochthonous tumor models validated the latter findings in an immunocompetent setting showing reduced tumor onset, tumor burden and prolonged survival. Collectively, these data demonstrate that miRISC assembly represents not only a marker for active miRNA-mediated gene repression but also an essential determinant of tumor development and progression. This principle appears to extend across tumors derived from different tissues, underscoring the critical role of this post-transcriptional regulatory layer in human cancers.

We propose that aberrant cell cycle and mitotic regulation underlie the proliferative defects observed in cancer cells with impaired miRISC. A plausible explanation is that miRNA inhibition leads to aberrant persistence of cell cycle related mRNAs beyond their physiologic window of operation, thereby affecting cell cycle checkpoints decisions and mitotic fidelity. This view aligns with the observation that controlled mRNA decay is essential for the correct phasing of most mRNAs during the cell cycle^128^.

Consistent with this model, the inhibition of miRNA function— whether through the LESKRES AGO2 mutations or direct interference of miRISC assembly—resulted in the selective aberrant de-repression of miRNA target genes involved in cell cycle regulation and mitotic division. We propose that this culminates in a failure to resolve chromosomal mis-segregation, resulting in chromosomal instability. These errors likely trigger DNA damage checkpoints, stalling cell cycle to facilitate the repair.

In support of this model, we provide evidence for increased G2/M checkpoint activation and extended mitotic duration (Figure 5G, S9A and S9B). Mitotic defects likely contributed also to the increased sensitivity to genotoxic agents we observed in miRISC impaired cancer cells (Figure 5A-5E).

Beyond cell cycle disruption, global mRNA stabilization upon miRISC inhibition can in principle interfere with transcriptome dynamics in response to metabolic or microenvironmental stress. This may further reduce tumor fitness, particularly in a dynamic *in vivo* setting^129^. Consistent with this line of reasoning are previous reports that miRISC function is largely dispensable for maintaining normal tissue homeostasis, yet becomes essential under conditions of acute stress and regenerative responses^94^.

Our data show that even transient miRISC inhibition (e.g., a single doxycycline pulse) can synergize with genotoxic therapies *in vitro*, suggesting that prolonged exposure may be unnecessary for therapeutic efficacy. These findings provide a rationale for developing AGO/miRISC-targeted therapies that exploit the unique dependency of malignancies on miRNA-mediated gene regulation.

## Limitations of the study

Because miRNAs regulate the expression dynamics of most mRNAs, global miRISC inhibition by T6B inevitably leads to pleiotropic effects. Therefore, although aberrant mitotic division appears to be a major driver of impaired proliferation and genotoxic sensitivity shared across various cancer models in our study, we cannot rule out the possibility that additional biological processes might be partially involved. Moreover, the tumor-specific transcriptomic landscape is likely to generate additional, context-dependent vulnerabilities in response to miRISC inhibition. For instance, T6B expression induced a gene signature indicative of suppressed KRAS signaling specifically in HCT116 cells, for which this pathway is a major oncogenic driver^130^. Future studies should systematically investigate cancer type-specific dysregulations—including critical genes, signaling nodes, and molecular mechanisms—to fully elucidate how miRISC inhibition shapes therapeutic responses across malignancies.

Our biochemical and computational analysis of cells and tissues that express T6B indicate that the peptide is capable of significantly disrupting miRISC function. However, it is likely that *in vivo* and *in vitro* some residual miRISC activity may persist. This scenario is even more likely in tumor regions with limited treatment access (e.g. doxycycline), leading to sub-optimal T6B expression. In some settings, this limitation could result in underestimating the actual extent to which impaired miRISC functions influence cancer development and progression.

## Acknowledgements

This work was supported by the NIH-NCI grant R01CA245507 (to A.V. and G.L.R.), by Deutsche Forschungsgemeinschaft (Le4223/4-1 to D.L) and FWF – Österreichischer Wissenschaftsfonds (I 6657-B to D.L.). We acknowledge the use of the following core facilities at the Memorial Sloan Kettering Cancer Center (MSKCC): The Molecular Cytology Core; The Mouse Genetics Core Facility; The Laboratory of Comparative Pathology, and the Integrated Genomics Operation Core, funded by the NCI Cancer Center Support Grant (CCSG, P30 CA08748), Cycle for Survival, and the Marie-Josée and Henry R. Kravis Center for Molecular Oncology. M.W. is supported by the Erwin Schrodinger Fellowship from the Austrian Science Fund (FWF, J4723). M.Z. is a recipient of the Paul Calabresi Career Development Award for Clinical Oncology from the Memorial Sloan Kettering K12 Clinical Translational Cancer Research Training Program. We thank Amaia Lujambio for donating the DNA vectors necessary to induce HCCs used in the present study, and members of her lab for technical guidance. We thank members of the Benezra laboratory for discussion and suggestions.

## Author contribution

M.Z., Z.J., G.L.R and A.V. designed the research project; Investigation: M.Z., Z.J., Y.C.F., I.L., X.L., M.A.Y., M.W., T.D., K.C.F., V.K.R., O.P., J.A.P., D.P., T.M., J.I.D and G.L.R.; Formal Analysis and Software: M.Z., Z.J., X.L., J.A.V. and A.V.; Writing of the original draft: M.Z., Z.J., D.L., A.V. and G.L.R with input from all coauthors.; Funding Acquisition: M.Z., D.L., A.V. and GLR; Resources: M.O., K.C. and C.M.; Supervision: M.Z., F.E.D., V.C., C.D.M., M.B.T., K.M.H., J.C., R.B., J.A.V., D.L., G.L.R and A.V.

**The authors declare no competing interests**.

## Materials and methods

### Animal husbandry

All animal experiments were conducted in accordance with protocols approved by the MSKCC Institutional Animal Care and Use Committee (IACUC) (protocol numbers: 10-10-022). Mice were housed in specific pathogen-free and maintained under a 12-hour light/dark cycle with ad libitum access to food. For experiments requiring doxycycline, the regular diet was replaced with irradiated doxycycline 625 ppm (blue) pellets (Test Diet). Endpoints were established based on appearance of signs of severe distress, as advised by the in-house veterinarian service.

### Animal models

To generate KP lung adenocarcinomas, recombinant adeno-CRE (ViraQuest) was delivered by intratracheal injection into the lungs of 10-12-week-old female KRas^LSL-G12D/+^; Trp53^fl/fl^ mice. Tumors were harvested 3 months after infection. Normal lungs were obtained from age and sex-matched mice. KP sarcomas were induced in the gastrocnemius of 8–10-week-old KP-rtTA or KP-T6B mice following in situ Adeno-CRE injection. Adeno-CRE was mixed with Matrigel® matrix (Corning, 354277) in a 1:1 ratio, and 50µL of the solution were used to inject each animal. Tumors were harvested at endpoint. To generate Eml4/Alk lung adenocarcinomas, 10-12-week-old mice were infected by intratracheal delivery of a recombinant adenovirus expressing Cas9 and the two sgRNAs necessary to induce the Eml4-Alk rearrangement, following a protocol previously described by Maddalo et al.^99^. Recombinant adenovirus used to induce the desired chromosomal rearrangements was purchased from ViraQuest. The B6.Cg-Tg(IghMyc)22Bri/J strain (Jax Strain #:002728) used to generate Eµ-Myc B cell lymphomas has been previously described by Adams et al.^98^. Eµ-Myc mice express a c-Myc transgene under the control of the B-cell-specific Eμ enhancer and develop B-cell lymphomas within 4–6 months of age. When mice showed signs of sever distress they were euthanized and tumors collected. To generate hepatocellular carcinomas, a mix of plasmids was injected into 4-6-week-old female mice by hydrodynamic tail vein injection as previously described by Molina-Sanchez et al. ^110^. A DNA mix containing 12.5µg MYC-IRES-Luc (Addgene #129775), 12.5µg px300-p53 (Addgene # 59910), and 2.5µg SB-13 (a gift from Amaia Lujambio) was prepared in sterile saline solution and injected in each animal. Mice were prewarmed under heat source for 5-10 minutes. Mice were injected with prepared DNA solution in a volume equivalent to 10% of the animal’s body weight using a ⅝-inch 26g needle in 5-8 seconds. The miR-92a knock-out mouse model was previously described in Han et al.^95^. Spleens of 8–10-week-old females were removed for B cell isolation and ex vivo activation. The R26^rtTA^; Col1a1^T6B^ (R26^T6B^) and R26^rtTA^; Col1a1^wt^ (R26^rtTA^) strains (Figure S3) have been previously described in La Rocca et al.^94^. To achieve expression of the T6B transgene in R26^rtTA^; Col1a1^T6B^ mice, a diet containing doxycycline (Test Diet 625 ppm pellets) was administered at variable time points according to the experimental design. The Col1a1^T6B^ strains is deposited at the Jackson Laboratory (JAX stock #036470).

### Xenograft models

Athymic nude mice (Jackson Laboratory) were injected subcutaneously with 0.5 million cells suspended in 100μl of 50% Matrigel (Corning, 354277) into the lower right and left dorsal flanks using a 27G needle to induce tumor xenografts. Tumor volume was measured twice a week using a caliper to determine both length and width. Tumor volume was calculated using the formula: V = (short diameter × short diameter × long diameter)/2, approximating the tumor as an ellipse. In vivo YFP expression was captured using the IVIS Spectrum Imaging System (PerkinElmer) and analyzed with Living Image® software. Mice were anesthetized with 2.5-3.5% isoflurane in air during imaging. Images were taken from both flanks, with parameters adjusted to avoid signal saturation and consistently maintained across all mice.

### Lung tumor burden measurement

Computed tomography (µCT) of the entire lung for each animal was obtained at 2 months (baseline) and 3 months following intratracheal delivery of adenovirus as described above. The entire lung from apex to base was then manually segmented on each slice using ImageJ, and the mean density was used to define lung tumor burden. Lung density of each mouse was first normalized to the liver density of the same scan, and then the liver-normalized lung density at 3 months was then secondarily normalized to the liver-normalized lung density at 2 months for the same animal to measure relative tumor burden.

### *In vivo* radiation of the hindlimb

Hind limb irradiation was performed on mice with a palpable tumor-bearing left hind limb when the tumor reached at least 100mm^3^ as measured by calipers using the formula V = (pi*x*y*z)/6, where V defines volume, and x, y, and z define length, width, and height of the tumor. At the time of radiation delivery, mice were restrained and shielded with commercial partial body radiation shields (Braintree, MHS1-F LF) and placed on the irradiation stage in the X-RAD 320 (Precision X-ray). Radiation was delivered with an open field with restrained mice shielded with exception of the exposed left hind limb. Mice were then followed for tumor progression as defined by tumor growth past quadrupling the initial volume on the day of the treatment. We used tumor-quadrupling as definition of tumor progression in this context, as transient local tissue swelling immediately after radiation can lead at least “tumor tripling” by caliper measurement, which subsequently resolves several days after treatment.

### Tissue isolation

Mice were euthanized via CO_2_ overdose before tissues isolation. Tumors and relevant organs were extracted from 8-to 15-week-old mice, washed with PBS, snap-frozen in liquid nitrogen and stored at −80°C until further processing. A portion of the tissues was fixed in 10% neutral buffered formalin and embedded in paraffin in a Leica ASP6025 tissue processor for histopathology and immunohistochemistry analyses. A portion of tumor samples was digested and cultured to generate cell lines. More specifically, tumors were first mechanically dissociated using scalpels, then enzymatically digested in a solution of 200 µg/ mL Collagenase IV and 100 µg/mL DNase I in PBS at 37°C for 1 hour, with gentle agitation every 10 minutes to facilitate tissue breakdown. After digestion, samples were centrifuged at 200g for 5 minutes, and the resulting cell pellets were washed three times with DMEM/F12 medium containing 1% penicillin-streptomycin and 1% GlutaMAX, then filtered using 70uM cell strainers. The derived tumor cells were then cultured in DMEM supplemented with 10% FBS, 1% penicillin-streptomycin, and 1% GlutaMAX for downstream applications.

### B cells harvesting and activation

Splenic B cells were harvested and dissociated into single-cell suspensions by mashing the spleen through a 70 μm strainer using the back end of a 1 mL syringe plunger. Naïve B cells were then isolated by negative selection according to the manufacturer’s instructions using anti-CD43 microbeads (Miltenyi Biotec). Three million naïve B cells were seeded at a concentration of 1 x 10^6^cells/mL in a six-well non-treated culture dish in B cell media (RPMI 1640 with L-glutamine (Gibco) supplemented with 15% fetal bovine serum (Corning), 1% penicillin-streptomycin (GeminiBio), 2 mM L-glutamine (Memorial Sloan Kettering Cancer Center Media Preparation Facility), and 55 μM βMercaptoethanol (Gibco)). B cells were then cultured in B cell media supplemented with 33 μg/ml LPS plus 25 ng/ml IL-4 (R&D Systems) for 48 hours at 37°C in a humidified 5% CO_2_ incubator.

### Cell lines and cell culture conditions

T6B- and CTL-expressing cell lines were generated by retroviral transduction of a panel of parental human and mouse cell lines. Human parental lines HCT116 (#CCL-247), HeLa (#CCL-2), A549 (#CCL-185) were purchased from ATCC. Mouse parental KP lung adenocarcinoma, KP sarcoma, and hepatocellular carcinoma cell lines were derived from the corresponding tumors generated in mice as described in section “Animal models”. The human osteosarcoma cell line was derived from ta metastasis found in the right lower lobe of the lung of an osteosarcoma patient according to our MSKCC IRB approved protocols (MSKCC-IRBs:06-107 and 12-245). Cell lines were maintained in log-phase growth in a humidified incubator at 37°C, 5% CO2 prior to experimental manipulation and were grown in Dulbecco’s Modified Eagle Medium (DMEM) supplemented with 10% heat-inactivated FBS, 10 U/ml penicillin/streptomycin, and 2 mM GlutaMAX, unless otherwise specified. HCT116 cells were maintained in McCoy’s medium supplemented with 10% heat-inactivated FBS, 10 U/ml pen/strep, and 2 mM L-glutamine. PDX osteosarcoma cells were maintained in RPMI 1640 Medium supplemented with 10% heat-inactivated FBS, 10 U/ml pen/strep, and 2 mM GlutaMAX (Gibco # 35050061).

### Retroviral particle production and cell line transduction

Sequences coding for the YFP- and BFP-tagged version of the T6B and CTL fragments (Table 1), where subcloned in a modified version^94^ of the retroviral expression vector pSIN-TREtight-HA-UbiC-rtTA3-IRES-Hygro (Addgene #20734), an all-in-one Tet-on vector that allows constitutive expression of the reverse tetracycline-regulated trans-activator (rtTA3) gene, and doxycycline-dependent expression of the transgene of interest. Cloning was carried out by using standard subcloning protocols. The sequence of each construct was confirmed by whole plasmid sequencing by Plasmidsaurus using Oxford Nanopore Technology with custom analysis and annotation. To generate retroviral particles, retroviral vectors were co-transfected along with packaging plasmids pCL-Eco (Addgene #12371) and VSV-G (Addgene #14888) into HEK-293T. Medium containing retroviral particles was collected after 48h and filtered using a syringe filter unit with a 0.45 µm cut-off. Transduction of cell lines of interest was performed using harvested viral medium supplemented with polybrene 0.2 µl/ml, (#TR-1003G, Millipore Sigma).

### Growth curves

Cells were maintained in log-growth phase in appropriate media and seeded onto 6-well plate at 50,000 cells per well. The next day, cells were counted at indicated time points using Cytation C10 Confocal Imaging Reader (BioTek) using YFP, BFP, or bright field, as applicable to each cell type and experimental design. For each time point and cell type, 3 wells were used for analysis. All growth curves were normalized to their respective 0-hour values.

### Competitive cell growth assay

T6B-YFP and CTL-BFP cells of each cell type were maintained separately in log-growth phase in appropriate media with doxycycline for 24 hours to induce respective peptide expression. The next day, T6B and CTL cells were mixed in a 1:1 ratio, confirmed by equal number of YFP and BFP positive cells using flow cytometry. The cell mixture was seeded onto a 6-well plate at 200,000 total cells per well. Changes in percentages of YFP and BFP positive cells were measured at indicated timepoints using flow cytometry. For experiments involving doxorubicin, doxorubicin at indicated concentration for each cell type is added into cell mixture during seeding and maintained through the course of the experiment. Competition assays with and without doxorubicin were always performed in parallel using the same initial cell mixture.

### Quantitative cell cycle analysis

Cell cycle analysis was performed using dual labeling of DAPI and Edu, using Click-iT™ Plus EdU Alexa Fluor™ 647 Flow Cytometry Assay Kit (Invitrogen, #C10634) according to the manufacturer’s instructions. Briefly, cells in log-growth phase are incubated with 10µM Edu for 1h, then collected, fixed and permeabilized using buffers provided by the kit, and stained for DNA content using DAPI before being analyzed by flow cytometry. DAPI separates cells in G1 from those in G2/M, and Edu labels cells in S phase.

### Flow cytometry

Trypsinized cells are washed once with PBS and collected by centrifugation at 300× g for 5 minutes. Cells are then resuspended in FACS buffer (PBS with 2% FBS) and filtered through a 70 µm cell strainer to create single-cell suspension. Samples are analyzed on a LSR Fortessa Cell Analyzer (BD Biosciences). For each parental cell type, the same gatings are applied throughout experiments. Single color samples are used as compensation controls when appropriate.

### Clonogenic assays

For the radiation clonogenic assay, cells were plated in low density in triplicates, where cell number is directly correlated to radiation dose to account for increased cell killing and is empirically determined before the experiment by testing different cell numbers per dose. Cells were allowed to attach to the plate prior to addition of doxycycline at 1µg/ml in the media 24 hours prior to irradiation. After irradiation, surviving cells were allowed to grow for up to 1 weeks on doxycycline-free medium to determine clonogenic survival. Colonies were fixed with 70% ethanol, stained with 0.01% methylene blue and counted under a dissecting microscope. Only those with at least 50 cells were included in the final tabulation. Raw percent survival is determined by the number of counted colonies divided by the number of cells plated. Raw percent survival at each dose point is then normalized to that at 0 Gy for the radiation clonogenic assay.

### Immunofluorescent staining for assessment of mitotic errors

HCT116 cells were treated with 1µg/ml doxycycline for 3 days prior to seeding in six well plate on poly-(L)-Lysine coated coverslips. After allowing cells to reattach for 24h while keeping them in doxycycline supplemented media, cells were then fixed with 4% paraformaldehyde. Primary human fibroblasts were allowed to reach near confluence and were seeded directly on coverslips 24h prior to fixation at 30% confluency. After fixation, cells were permeabilized, and counterstained. For this, following primary antibodies were applied diluted in 5% BSA in PBS in indicated dilution ratio: anti-CREST (Proteintech; rabbit, 1:50), anti-β-catenin (BD biosciences; mouse, 1:100), anti-tubulin (mouse, 1:100), and anti-pericentrin (rabbit, 1:500). Afterwards, cells were incubated in secondary antibodies solutions: AlexaFluor488 (ThermoFischer; anti-mouse, 1:500), AlexaFluor594 (ThermoFischer; anti-mouse, 1:500) and AlexaFluor645 (ThermoFischer; anti-rabbit, 1:500). Cells were imaged the next day on BZ-X800 (KEYENCE) or on Fluoview FV3000 (Olympus). Analysis software used to count cells was either BZ-X800 Analyzer (Keyence) or cellSens (EVIDENT).

### Micronuclei assay

Cells were plated in 4 well chamber slides and exposed to doxycycline as described for radiation clonogenic assay. Cells were then allowed to grow for an additional 24 hours following exposure to 0 or 8 Gy and were then fixed with 4% paraformaldehyde and counterstained with DAPI. Images were taken at 20x, and total number of nuclei and number of nuclei with associated micronuclei or nuclear fragmentation were counted In ImageJ. A minimum of 311 nuclei were counted per clone per condition across 4 isogenic clones expressing either T6B or CTL peptide.

### G2/M boundary synchronization

To synchronize cells at the G2/M boundary, cells in log-growth phase are first synchronized to the G1/S boundary by treatment with thymidine (2.5 mM) for 16 hours. Cells are then released into drug-free media for 6 hours to progress through S phase, followed by a 6-hour treatment with nocodazole (100 ng/ mL) to arrest them at G2/M boundary. After two PBS washes, cells are released into drug-free media and allowed to proceed through mitosis. For downstream flow cytometry and western blot analyses, cells are harvested at fixed time intervals on ice using cell scrapers.

### Tissue and cell whole cell lysates

After harvesting, tissues are pulverized using a mortar, resuspended in 1 ml of lysis buffer per cm^3^ of tissue, and dounce-homogenized with a tight pestle. Total lysates of homogenized tissue are then cleared by two consecutive steps of centrifugation at 20,000× g for 5 minutes each. Two different lysis buffers were used, depending on specific downstream application. For size-exclusion chromatography (SEC), lysates were prepared in SEC buffer (150 mM NaCl, 10 mM Tris-HCl pH 7.5, 2.5 mM MgCl2, 0.01% Triton X-100). For western blotting, lysates were prepared in RIPA buffer (Santa Cruz Biotechnology #24948). Both buffers were supplemented with EDTA-free complete protease inhibitors (Sigma-Aldrich #11836170001), phosphate inhibitors (Roche #04906837001), and 1 mM DTT upon use. To prepare total extracts from cultured cells, pelleted cells are snap frozen in liquid nitrogen and stored at −80°C until further processing. Pellets are then resuspended in lysis buffer and incubated for 10 min on ice. Total lysates of homogenized tissue are then cleared by two consecutive steps of centrifugation at 20,000× g for 5 minutes each.

### Size-exclusion chromatography

Size-exclusion chromatography (SEC) was performed using Superose 6 10/300 GL prepacked column (GE Healthcare) equilibrated with SEC buffer (150 mM NaCl, 10 mM Tris-HCl pH 7.5, 2.5 mM MgCl2, 0.01% Triton X-100) as previously described^75^. Briefly, 400μl (1.5–2 mg) of cleared total lysates are run through the SEC column at a flow rate of 0.3 ml/min. 1-ml fractions are continuously collected. Proteins from each fraction are then extracted by TCA precipitation following standard protocol and run on SDS-PAGE gels for Western blotting analysis.

### Western blotting

All Western blotting experiments were performed using the Novex NuPAGE SDS/PAGE gel system (Invitrogen). Total tissue or cell lysates are run on either 3–8% Tris-acetate or 4–12% Bis-Tris precast gels, transferred to nitrocellulose membranes, and probed with antibodies specific to proteins of interest. Detection and quantification of blots are performed on either Amersham hyperfilm ECL (Cytiva #28906839) or Odyssey Imaging System (LICOR). The following commercial primary antibodies are used: anti-Ago2 (Cell Signaling #2897), anti-panAGO clone 2A8 (Sigma-Aldrich #MABE56), anti-GAPDH (Cell Signaling #2118), anti-alpha Tubulin (Cell Signaling #3873), anti-HA (Cell Signaling #C29F4), anti-Caspase 3 (Cell Signaling #9662), anti-Cleaved Caspase 3 (Cell Signaling #9661), and anti-p16/ INK4A (Cell Signaling #92803). Radixin used as lading control in Figure S1D was detected with anti-panAGO antibody^131^. The following commercial secondary antibodies are used: IRDye 800 anti-Rabbit (#926-32213, LICOR), IRDye 680 anti-Mouse (#926-68072, LICOR), anti-rabbit IgG, HRP-conjugated (GE Healthcare #NA934), and anti-mouse IgG, HRP-conjugated (GE Healthcare #NA931). Densitometric quantification of blots in Figure 5G was performed with ImageJ.

### Immunohistochemistry (IHC)

For immunohistochemistry, deparaffinized sections were subjected to antigen retrieval and processed with the EnVision + HRP kit (K401111-2, DAKO, Glostrup, Denmark) according to the manufacturer’s instructions. A primary polyclonal antibody against GFP at 1:400 dilution was diluted in Antibody Diluent (DAKO #S0809) and incubated overnight at 4°C. Next, sections were incubated in the provided anti-rabbit HRP-labeled polymer reagent, and detection was performed according to the manufacturer’s protocol. Images were acquired using an Olympus BX-UCB slide scanner.

### RNA sequencing

Total RNA from KP lung adenocarcinomas cells and B-lymphocytes were extracted using TRIzol Reagent (Invitrogen) according to the manufacturer’s instructions and then treated with DNase (QIAGEN). Extracted total RNA was quantified with RiboGreen assay (Invitrogen) and quality-controlled by BioAnalyzer (Agilent). 500 ng of total RNA with RIN values of 7.0–10 was subjected to polyA selection and TruSeq library preparation according to the instructions by Illumina (TruSeq Stranded mRNA LT Kit, Cat#RS-122-2102), with eight cycles of PCR. Paired-end RNA libraries were sequenced by the MSKCC Integrated Genomics Core. STAR v2.5.3a^132^ was used to align RNA reads to the standard mouse genome (mm10). Aligned reads were counted at each gene locus. DESeq2^133^. Moderated estimation of fold change and dispersion for RNA-seq data with DESeq2was used to perform differential gene expression analysis on expressed genes, and log2-fold changes were determined comparing T6B- and CTL-expressing KP LUAD cells, and between miR-92a KO and wild-type B-lymphocytes in resting and activated states, respectively. Small RNA quantification was performed by reanalyzing small RNA sequencing datasets published in Sala et al.^88^, which can be found at the Gene Expression Omnibus (GEO) (GSE203049).

## SUPPLEMENTAL FIGURES

**Figure S1.**
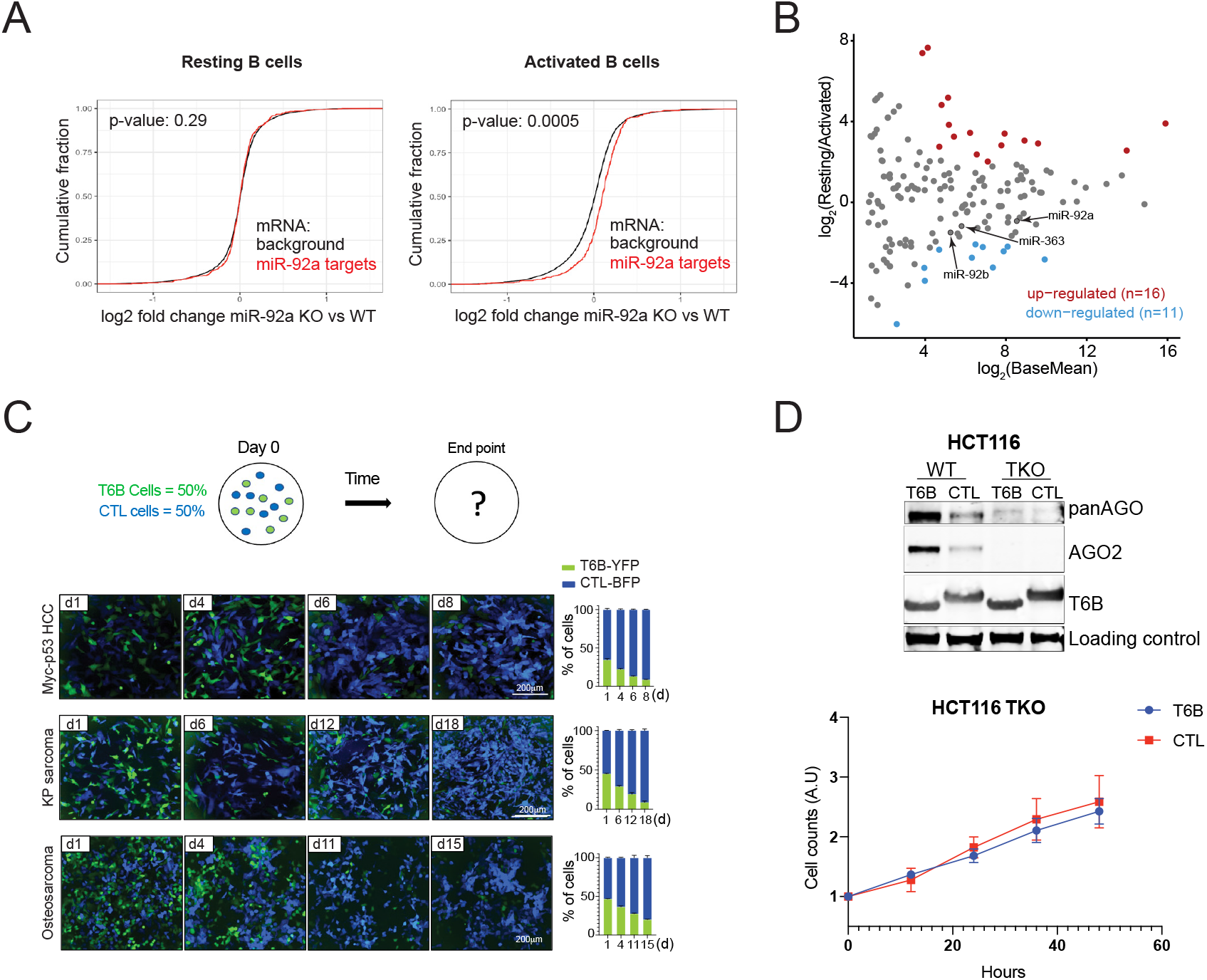
Inhibition of miRISC function impairs tumor growth. **(A)** Left, cumulative distribution plot of log2-fold changes in expression of predicted targets of miR-92a in miR-92a KO vs WT resting B cells. Right, cumulative distribution plot of log2-fold changes in expression of predicted targets of miR-92a in miR-92a KO vs WT activated B cells. Note that miR-92a targets (red lines) are preferentially upregulated compared to the general mRNA population (black lines) in activated B cells but not in resting B cells. **(B)** Scatter plot showing changes in expression levels of the indicated miRNAs upon B cell activation. miR-92a family members are labeled. **(C)** Representative epifluorescence images of cell lines derived from mouse hepatocellular carcinomas (HCC, top), mouse KP sarcoma (middle) and patient-derived osteosarcoma xenograft (bottom) at the indicated timepoints over the course of competitive cell growth assays (n = 3). Cells expressing CTL peptide tagged by BFP are blue. Cells expressing T6B peptide tagged by YFP are green. Right, corresponding quantifications of the percentages of BFP and YFP positive cells for each experiment. Error bars indicate mean ±SEM. **(D)** Top, Western blots showing the levels of expression of the indicated proteins in HCT116 AGO1,2,3 triple knock out (TKO). The antibody “panAGO” recognizes all AGO paralogues (AGO1, 2, 3 and 4, *∼*100 kDa). Radixin signal (*∼*70 kDa) has been used as loading control. Bottom, growth curves of HCT116 TKO upon expression of either T6B or CTL peptide.

**Figure S2.**
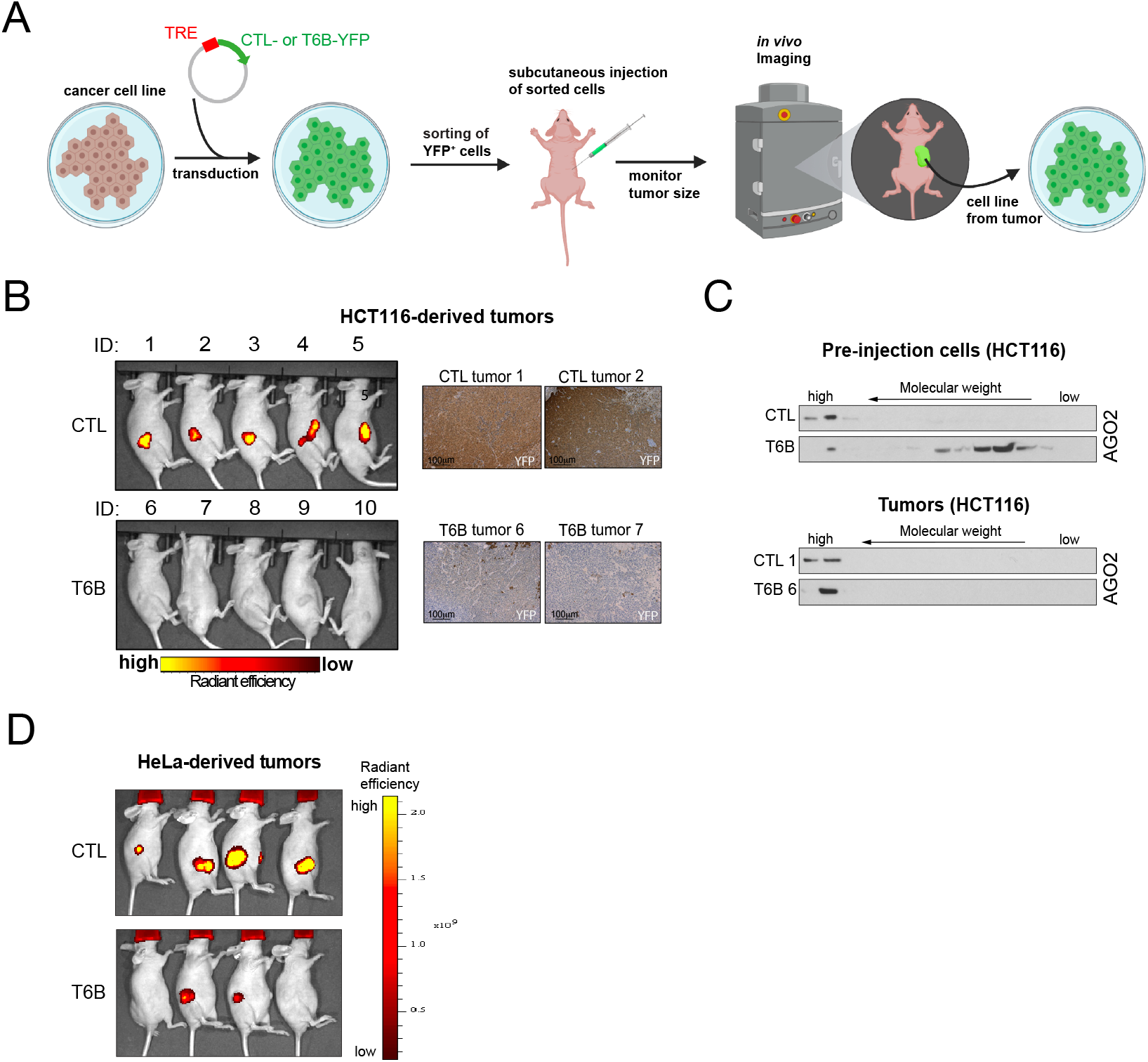
Selection against T6B expression in xenografts. **(A)** Schematics of experimental flow to generate and analyze xenografts and xenograft-derived cell lines. Both T6B and CTL peptides are tagged by YFP. **(B)** Left, scans taken by in vivo imaging showing YFP fluorescence intensity in T6B- or CTL-expressing tumors derived from HCT116 cells at endpoint. Right, immunohistochemistry (IHC) staining using an antibody against the HA-tag showing the expression levels of T6B and the CTL peptide in sections of the indicated formalin-fixed/paraffin-embedded tumors. **(C)** Western blots showing SEC profiles of AGO2 complexes in pre-injected HCT116 cells expressing either CTL or T6B peptide (top panels) and in cells derived from their tumors (bottom panels) at endpoint. **(D)** Scans taken by in vivo imaging showing YFP fluorescence intensity in T6B- or CTL-expressing tumors derived from HeLa cells at endpoint.

**Figure S3.**
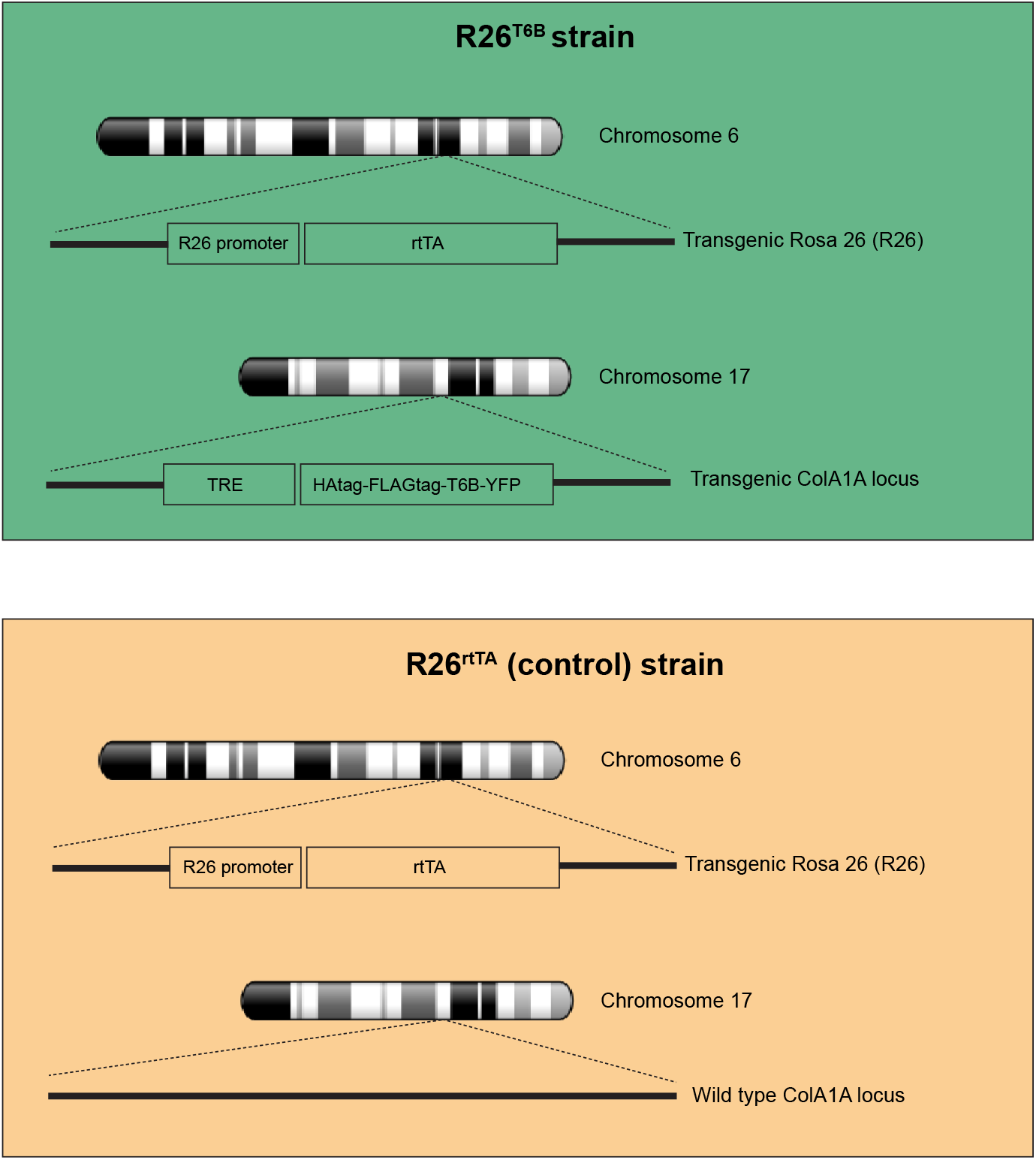
Schematics of the *R26*^*rtTA*^; *Col1a1*^*T6B*^ (R26^T6B^) and *R26*^*rtTA*^; *Col1a1*^*wt*^ (R26^rtTA^) mice design. Both the R26^T6B^ and R26^rtTA^ mice strains have the reverse tetracycline trans-activator (rtTA) coding sequence inserted into the *Rosa26* (R26) locus, resulting in constitutive rtTA expression driven by the endogenous R26 promoter. In addition, the R26^T6B^ strain **(top)** has the HAtag-FLAGtag-T6B-YFP fusion protein inserted into the *Col1A1* locus and driven by the tetracycline responsive element (TRE), resulting in T6B fusion protein expression upon doxycycline administration. The control R26^rtTA^ strain **(bottom)** carries a wild type *Col1A1* locus, therefore lacks T6B fusion protein expression even upon doxycycline administration.

**Figure S4.**
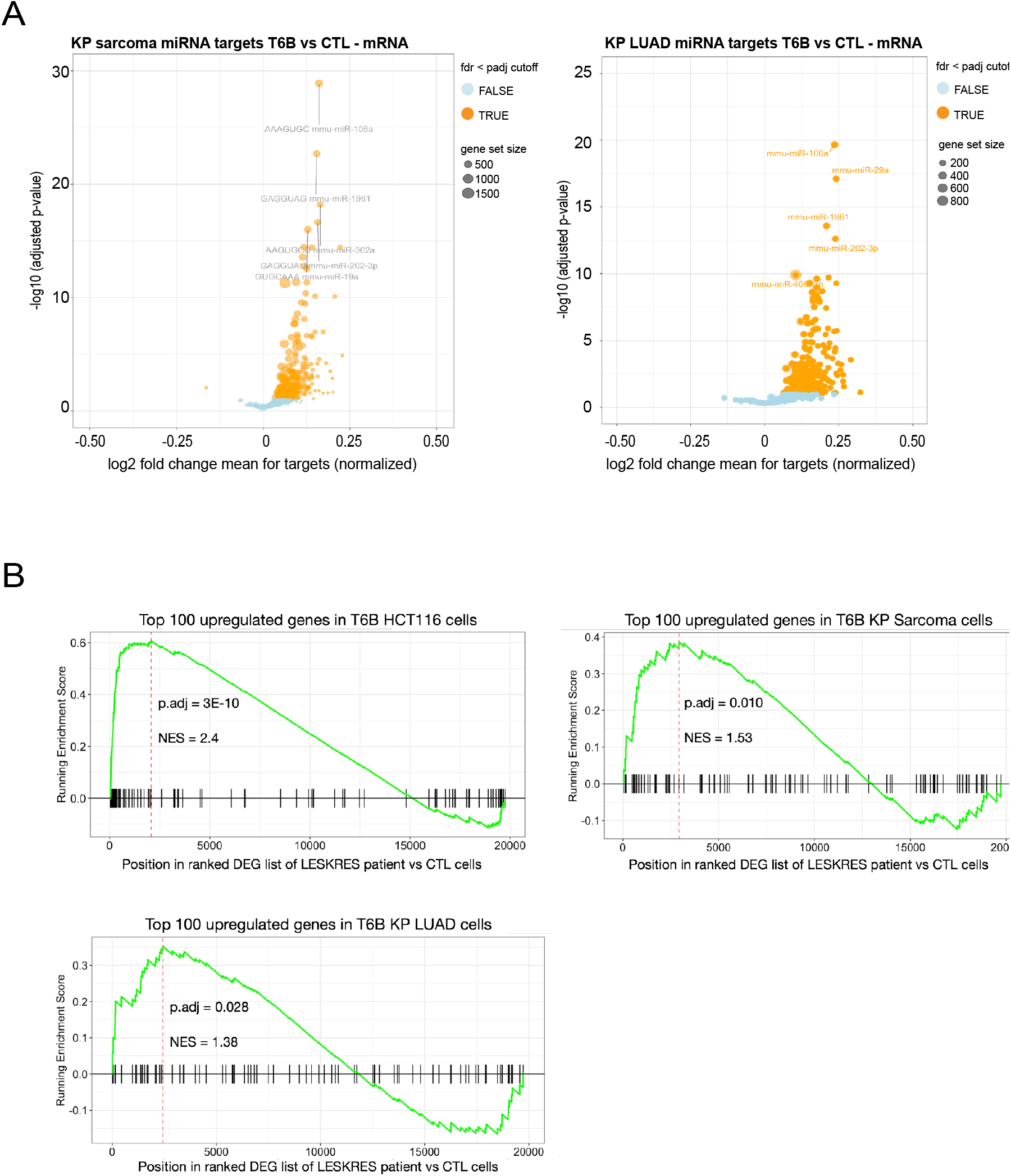
Transcriptome analysis in T6B-expressing cell lines. **(A)** Volcano plot of mRNAs showing preferential global up-regulation of predicted miRNA targets for each conserved miRNA family in KP sarcoma cells (left) and KP LUAD (right) expressing T6B compared to cells expressing the CTL peptide. Analysis on KP sarcoma cells was conducted after 72h hour doxycycline induction. Analysis on KP LUAD cells was conducted after 24h hour doxycycline induction. miRNA targets are grouped by circles, each representing a conserved miRNA family. The size of each circle is proportional to the number of predicted target genes within a given miRNA family. **(B)** Enrichment of top 100 most up-regulated genes in T6B-vs CTL-expressing cancer cells in the ranked list of differentially expressed genes in LESKRES patient-derived vs CTL fibroblasts. The Kolmogorov-Smirnov test was used to evaluate the significance of enrichment in (B).

**Figure S5.**
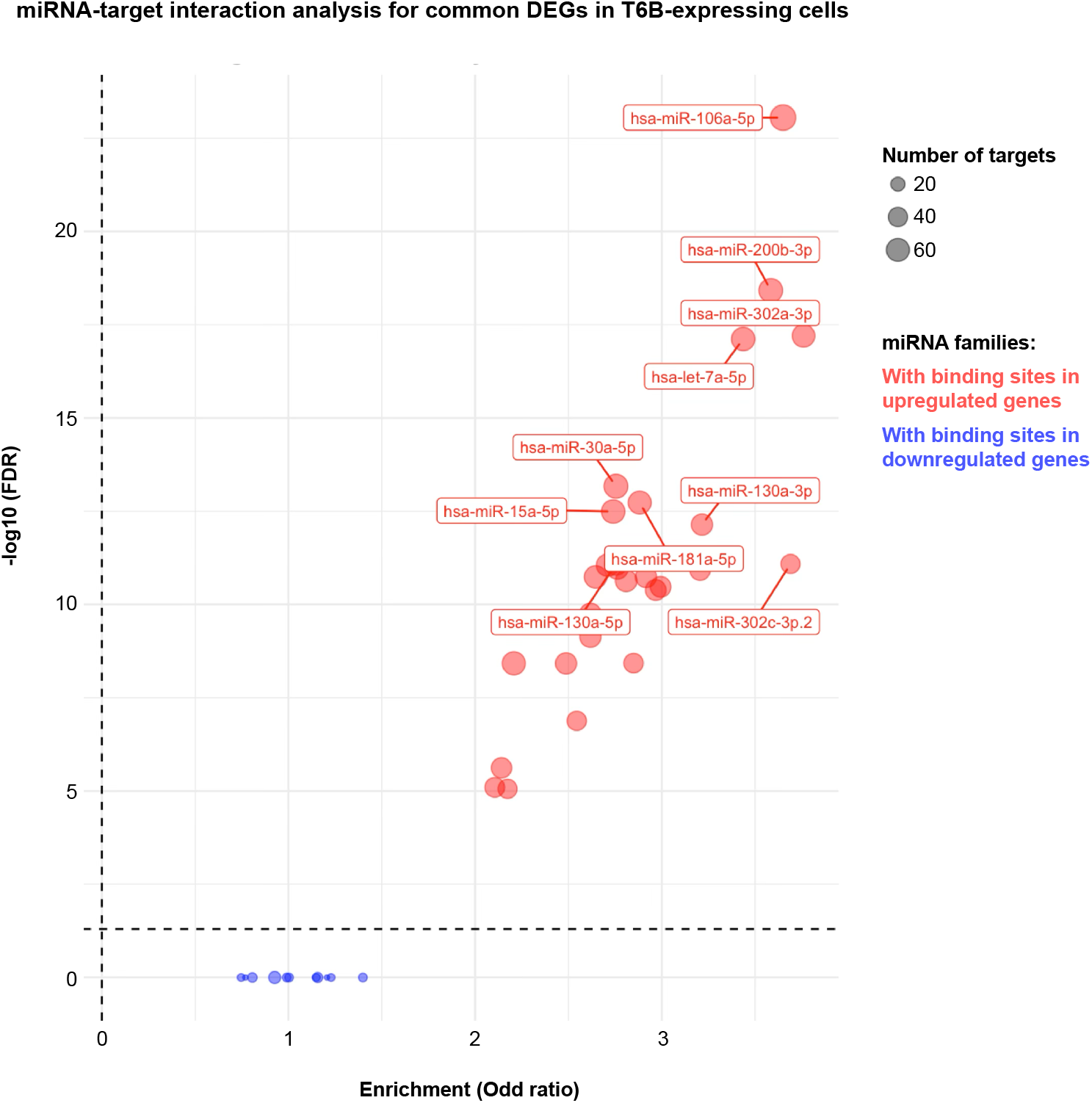
miRNA-target enrichment analysis for common differentially expressed genes (DEGs) in T6B tumor cells. Volcano plot of enriched miRNA families in commonly upregulated (red) and downregulated (blue) genes in T6B expressing tumor cell lines, identified by miRNA-target enrichment analysis tool MIENTURNET. The size of each circle is proportional to the number of target genes within a given miRNA family predicted by Targetscan. Top 10 most enriched miRNA families are shown. Note that only commonly regulated genes yield significantly enriched miRNA families.

**Figure S6.**
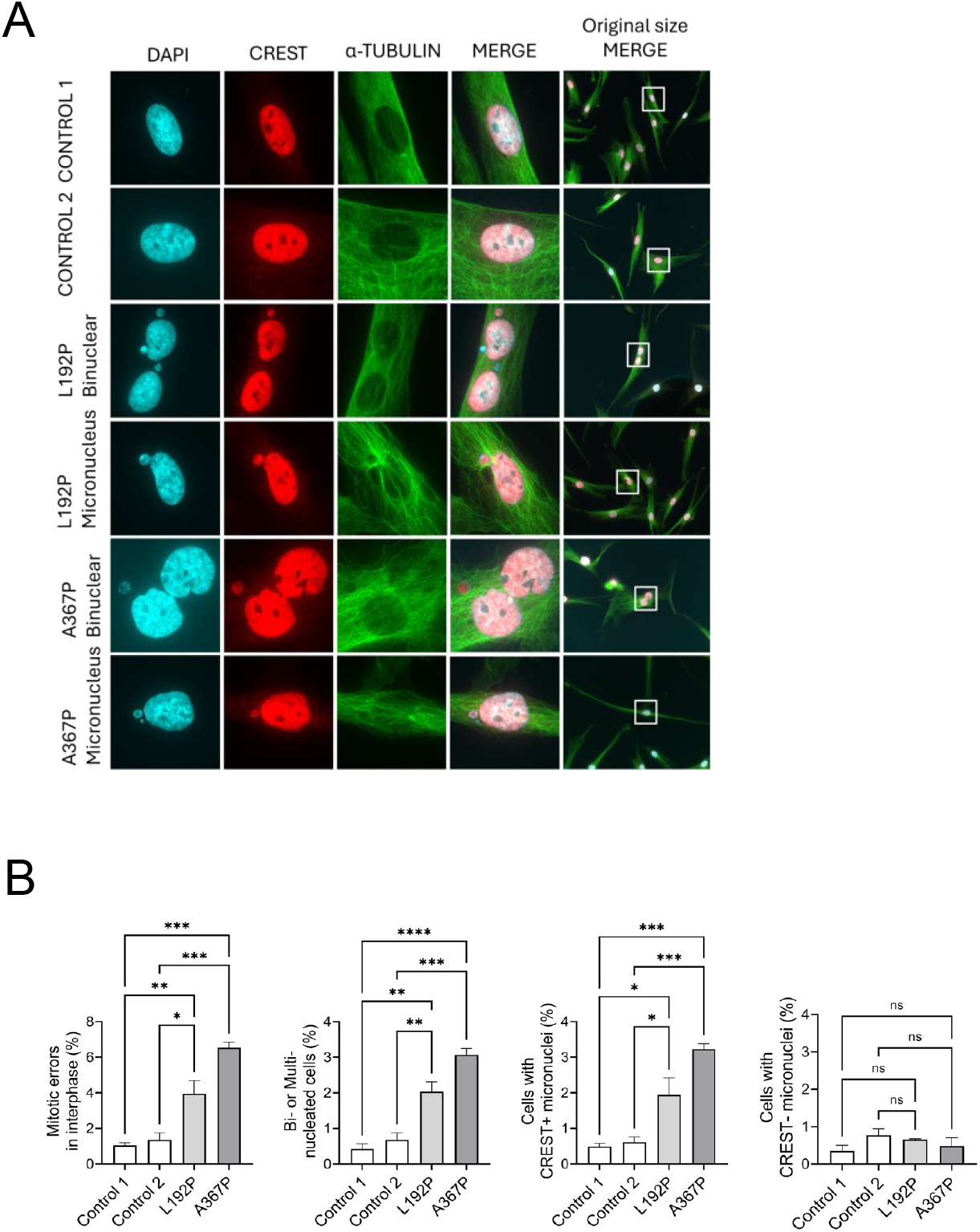
Cellular phenotype of LESKRES patient-derived fibroblasts. **(A)** Cellular phenotype of human primary fibroblasts from 2 healthy controls and two individuals with Lessel-Kreienkamp syndrome carrying specified changes in AGO2 protein (A367P and L192P). **(B)** Quantification of (A) with at least 200 cells analyzed per experiment. All quantifications are based on 3 independent experiments. Data are represented as mean ±SEM. * P<0.05, ** p<0.01, ***p<0.001, ****p<0.0001. Ordinary one-way ANOVA along the Tukey’s correction is used in (B).

**Figure S7.**
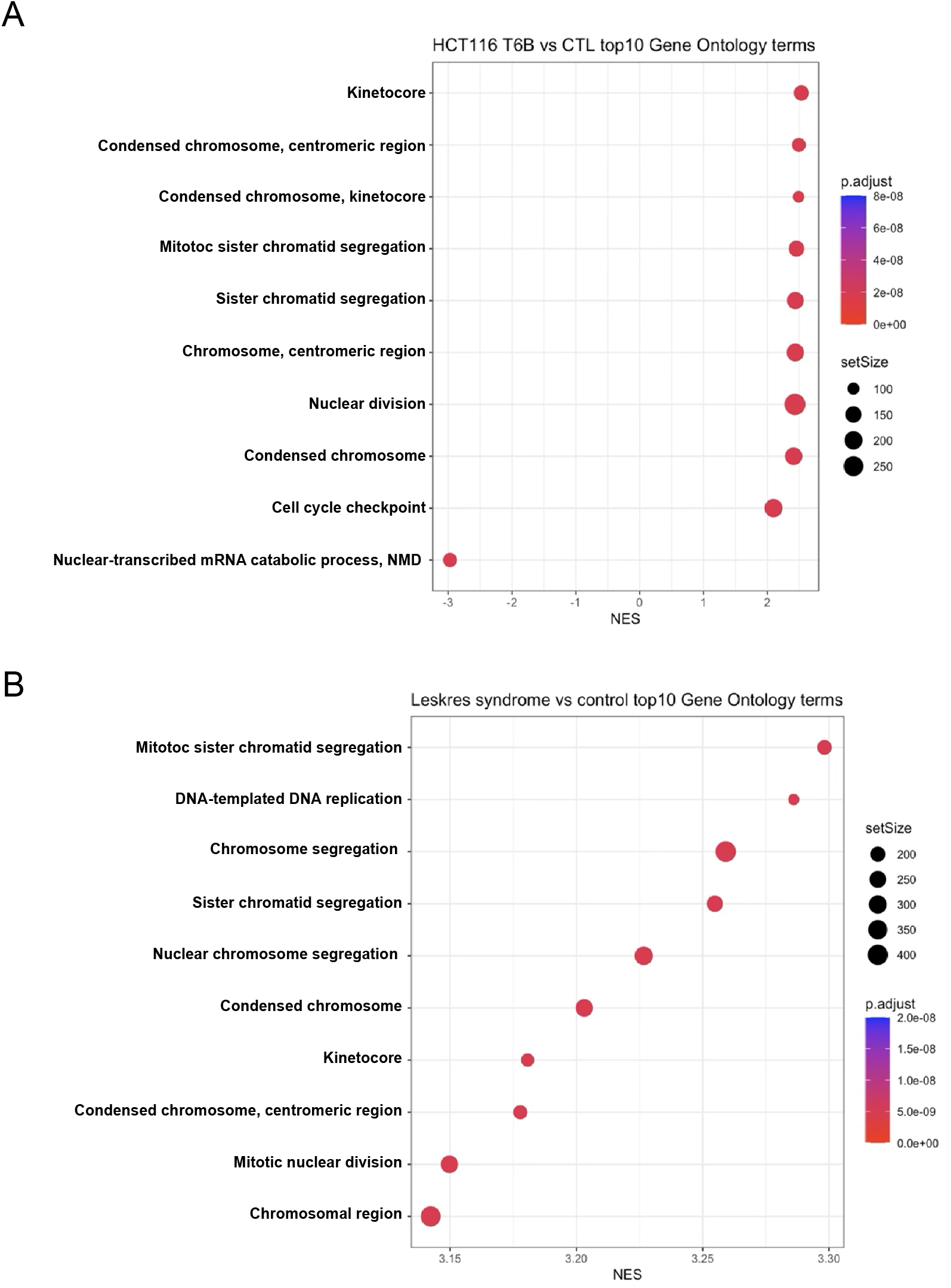
Gene Ontology Analysis in T6B-expressing HCT116 and LESKRES patient-derived fibroblasts. **(A)** Gene Set Enrichment Analysis (GSEA) of differentially expressed genes in HCT116 cells expressing T6B compared to CTL using annotated Gene Ontology (GO) gene sets. Top 10 differentially enriched terms are shown. **(B)** Same analysis as in (A) for LESKRES patient-derived fibroblasts compared to matched controls.

**Figure S8.**
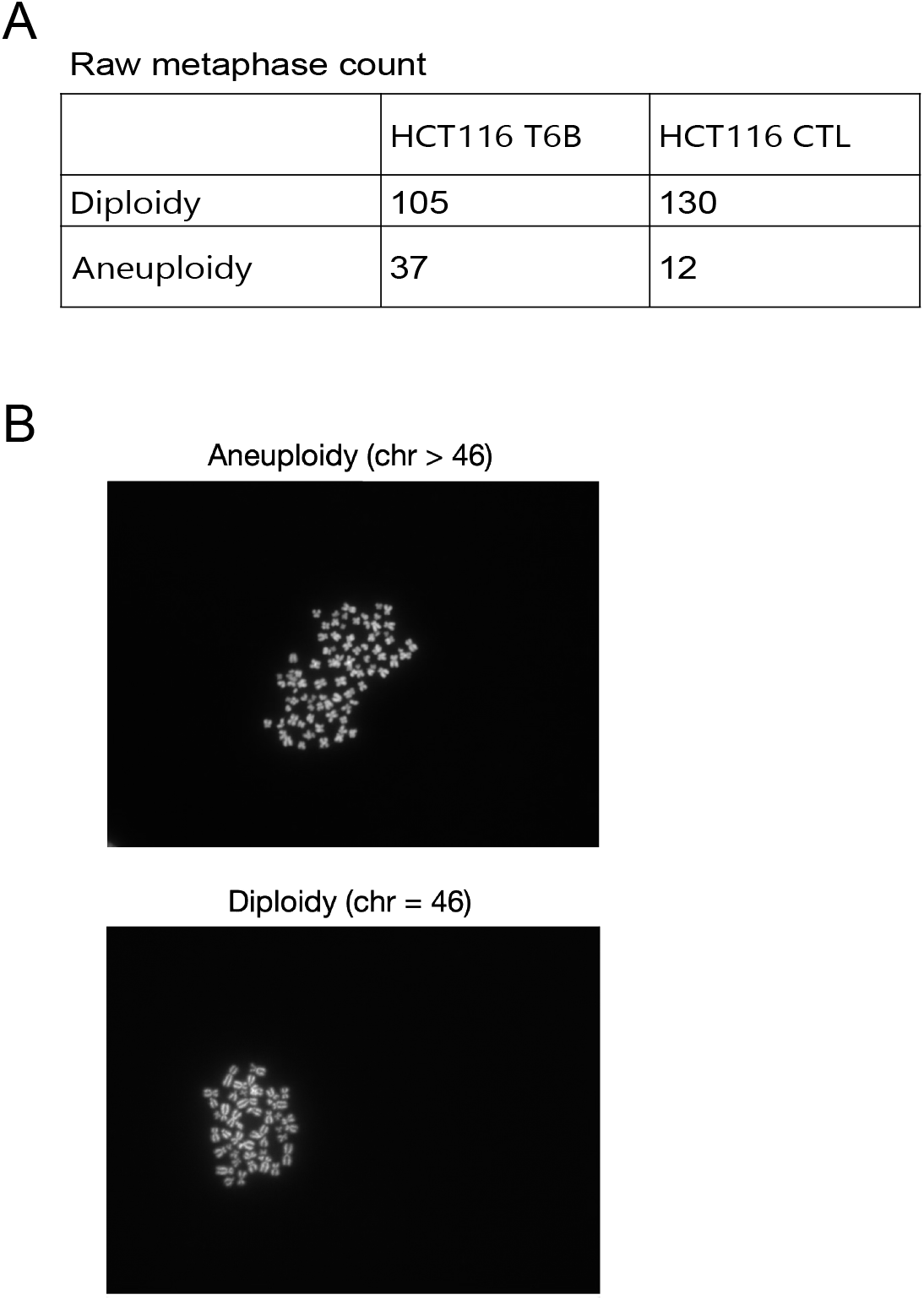
miRISC inhibition causes aneuploidy in cancer cells. **(A)** Left: Raw tabulations of counted metaphases of T6B- and CTL-expressing HCT116 cells harboring near/perfect diploidy or aneuploidy karyotype. **(B)** Representative images of metaphases displaying aneuploidy (top) and diploidy (bottom). Fisher’s exact test is used to analyze contingency in (A). ***p<0.001.

**Figure S9.**
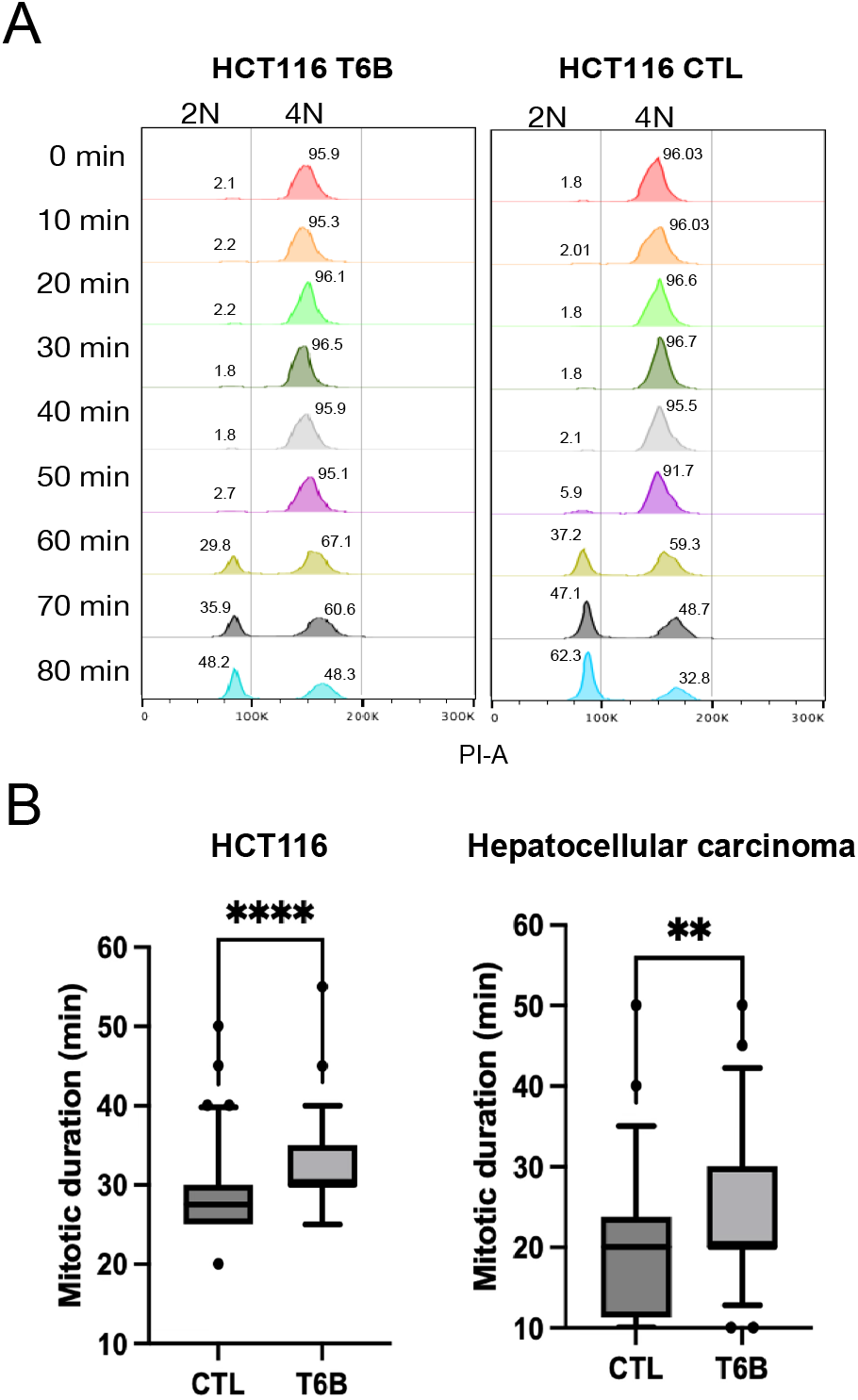
miRISC inhibition causes mitotic delays in cancer cells. **(A)** Actual flow cytometry plots related to Figure 5F showing DNA content in T6B and CTL HCT116 cells going through mitosis. **(B)** Mitotic duration measurement by phase contrast microscopy in HCT116 and hepatocellular carcinoma cells expressing T6B or CTL peptide. Two-sided Student’s t-test is used to compare matched T6B and control groups.

**Figure S10.**
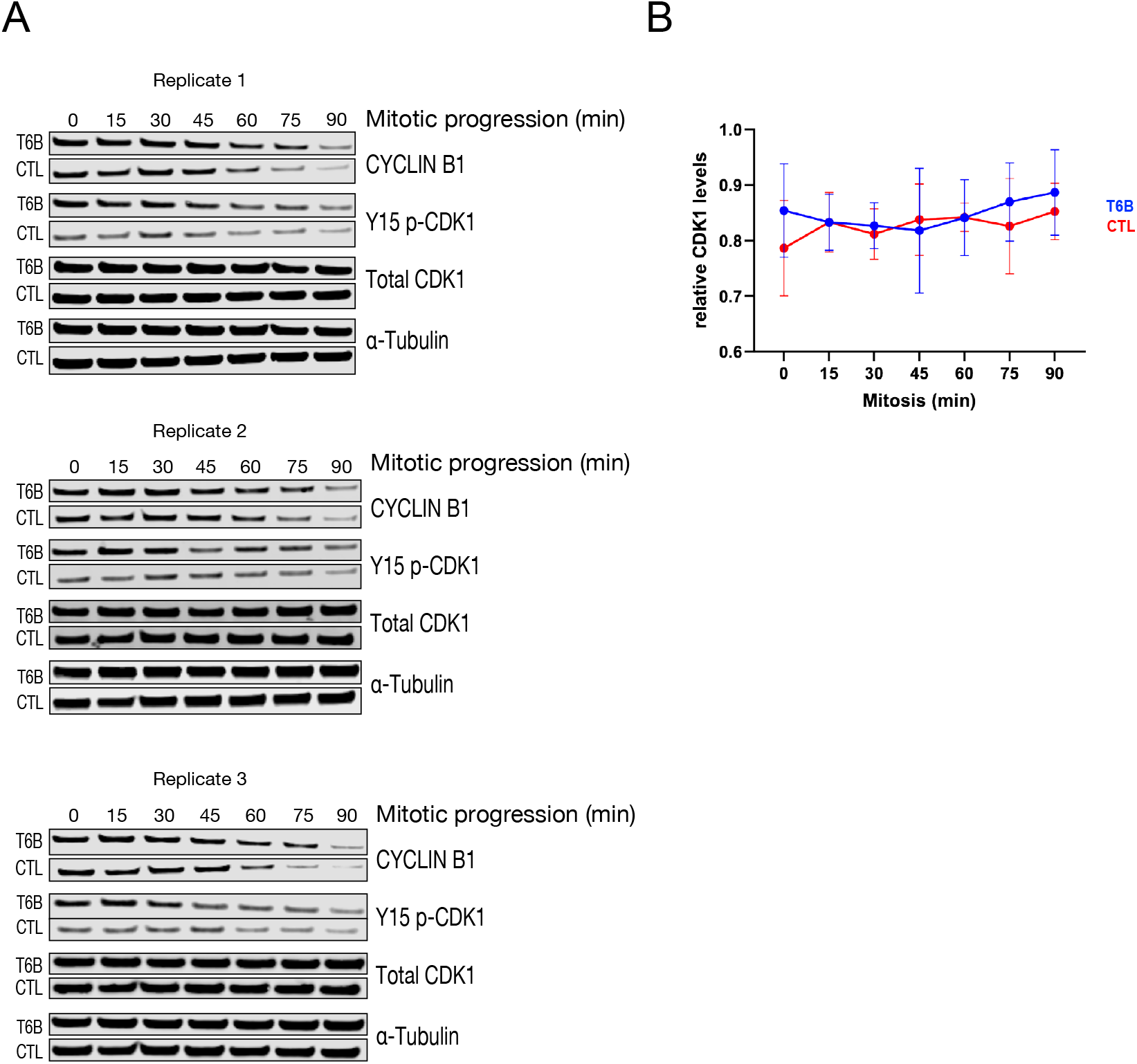
miRISC inhibition triggers G2/M checkpoint and delays CyclinB1 degradation. **(A)** Biological replicates of Western blot staining showing expression of the indicated proteins relative to plots showed in Figure 5G. **(B)** Quantification of signal of total CDK1 showed in (A).

**Figure S11.**
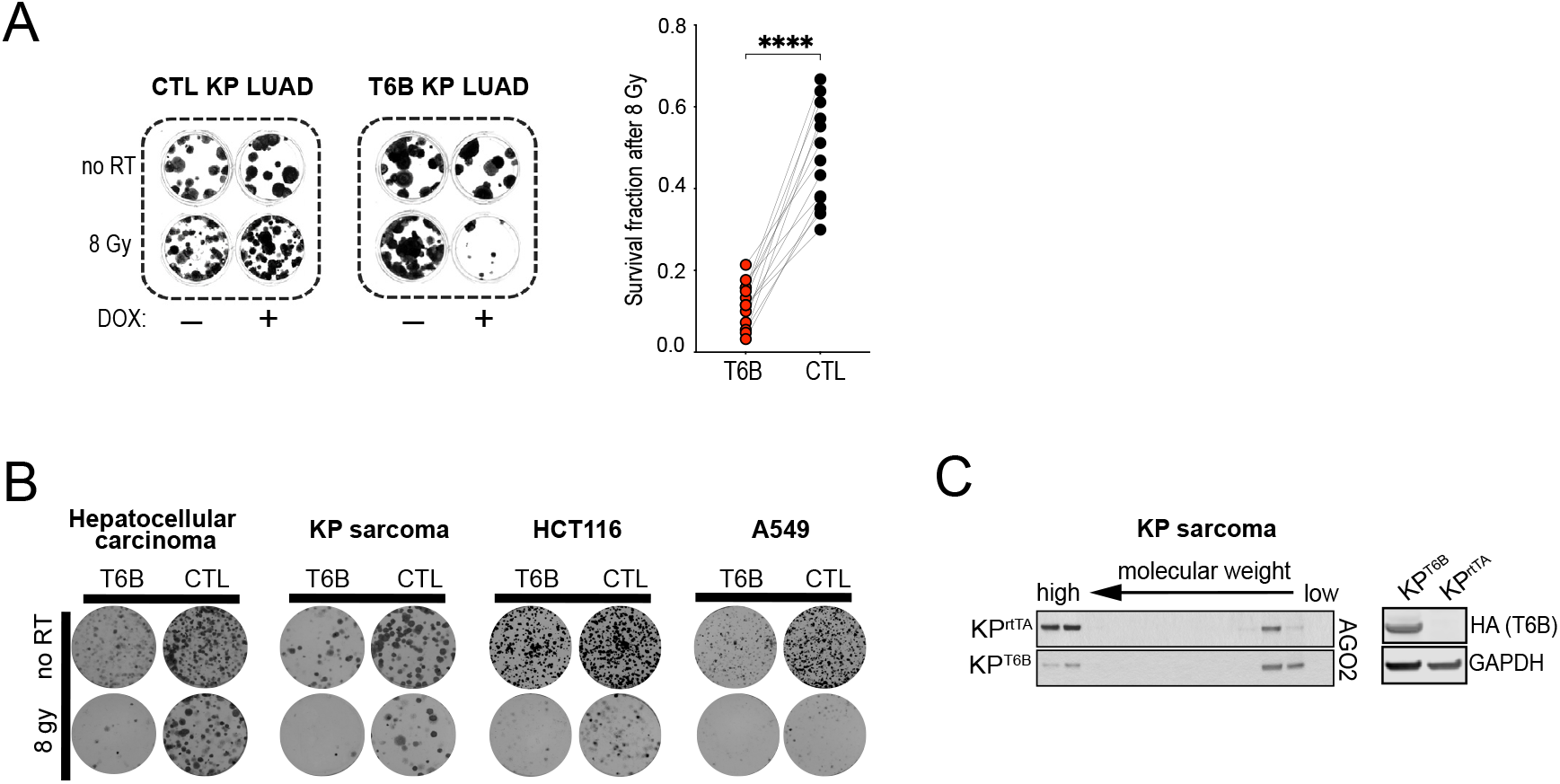
Inhibition of miRISC function sensitizes cancer cells to genotoxic agents. **(A)** Left: representative image of plates used in colony formation assays for CTL- and T6B-KP LUAD cells with or without pulsed-24-hour expression of corresponding fusion protein (dox+ and dox-, respectively), and with or without receiving a single dose of 8 Gy radiation (8 Gy and no RT, respectively). Right: Plots showing normalized values of colony counts of the assay shown in top left. **(B)** Representative image of plates used in colony formation assays for a variety of CTL- and T6B-expressing cells with pulsed-24-hour expression of corresponding fusion protein and with or without receiving a single dose of 8 Gy radiation (8 Gy and no RT, respectively). **(C)** Left, Western blots showing the AGO2 elution profile from SEC fractionation in KP-T6B and KP-rtTA sarcomas. Right, Western blots showing the level of expression of the T6B (detected using an anti-HA-tag antibody) induced in KP-T6B and KP-rtTA mice.

## References

1. Bartel, D.P. (2018). Metazoan MicroRNAs. Cell 173, 20–51. 10.1016/j.cell.2018.03.006.

2. Eichhorn, S.W., Guo, H., McGeary, S.E., Rodriguez-Mias, R.A., Shin, C., Baek, D., Hsu, S.H., Ghoshal, K., Villen, J., and Bartel, D.P. (2014). mRNA destabilization is the dominant effect of mammalian microRNAs by the time substantial repression ensues. Mol Cell 56, 104–115. 10.1016/j.molcel.2014.08.028.

3. Friedman, R.C., Farh, K.K., Burge, C.B., and Bartel, D.P. (2009). Most mammalian mRNAs are conserved targets of microRNAs. Genome Res 19, 92–105. 10.1101/gr.082701.108.

4. Kozomara, A., Birgaoanu, M., and Griffiths-Jones, S. (2019). miRBase: from microRNA sequences to function. Nucleic Acids Res 47, D155–D162. 10.1093/nar/gky1141.

5. Fromm, B., Hoye, E., Domanska, D., Zhong, X., Aparicio-Puerta, E., Ovchinnikov, V., Umu, S.U., Chabot, P.J., Kang, W., Aslanzadeh, M., et al. (2022). MirGeneDB 2.1: toward a complete sampling of all major animal phyla. Nucleic Acids Res 50, D204–D210. 10.1093/nar/gkab1101.

6. McGeary, S.E., Lin, K.S., Shi, C.Y., Pham, T.M., Bisaria, N., Kelley, G.M., and Bartel, D.P. (2019). The biochemical basis of microRNA targeting efficacy. Science 366. 10.1126/science.aav1741.

7. Flynt, A.S., and Lai, E.C. (2008). Biological principles of microRNA-mediated regulation: shared themes amid diversity. Nat Rev Genet 9, 831–842. 10.1038/nrg2455.

8. Treiber, T., Treiber, N., and Meister, G. (2019). Regulation of microRNA biogenesis and its crosstalk with other cellular pathways. Nat Rev Mol Cell Biol 20, 5–20. 10.1038/s41580-018-0059-1.

9. Mauro, M., Berretta, M., Palermo, G., Cavalieri, V., and La Rocca, G. (2023). The Multiplicity of Argonaute Complexes in Mammalian Cells. J Pharmacol Exp Ther 384, 1–9. 10.1124/jpet.122.001158.

10. Iwakawa, H.O., and Tomari, Y. (2022). Life of RISC: Formation, action, and degradation of RNA-induced silencing complex. Mol Cell 82, 30–43. 10.1016/j.molcel.2021.11.026.

11. Schirle, N.T., Sheu-Gruttadauria, J., and MacRae, I.J. (2014). Structural basis for microRNA targeting. Science 346, 608–613. 10.1126/science.1258040.

12. Braun, J.E., Huntzinger, E., Fauser, M., and Izaurralde, E. (2011). GW182 proteins directly recruit cytoplasmic deadenylase complexes to miRNA targets. Mol Cell 44, 120–133. 10.1016/j.molcel.2011.09.007.

13. Chekulaeva, M., Mathys, H., Zipprich, J.T., Attig, J., Colic, M., Parker, R., and Filipowicz, W. (2011). miRNA repression involves GW182-mediated recruitment of CCR4-NOT through conserved W-containing motifs. Nat Struct Mol Biol 18, 1218–U1262. 10.1038/nsmb.2166.

14. Chen, C.Y.A., Zheng, D.H., Xia, Z.F., and Shyu, A.B. (2009). Ago-TNRC6 triggers microRNA-mediated decay by promoting two deadenylation steps. Nat Struct Mol Biol 16, 1160–U1166. 10.1038/nsmb.1709.

15. Chen, Y., Boland, A., Kuzuoglu-Ozturk, D., Bawankar, P., Loh, B., Chang, C.T., Weichenrieder, O., and Izaurralde, E. (2014). A DDX6-CNOT1 complex and W-binding pockets in CNOT9 reveal direct links between miRNA target recognition and silencing. Mol Cell 54, 737–750. 10.1016/j.molcel.2014.03.034.

16. Fabian, M.R., Cieplak, M.K., Frank, F., Morita, M., Green, J., Srikumar, T., Nagar, B., Yamamoto, T., Raught, B., Duchaine, T.F., and Sonenberg, N. (2011). miRNA-mediated deadenylation is orchestrated by GW182 through two conserved motifs that interact with CCR4-NOT. Nat Struct Mol Biol 18, 1211–1217. 10.1038/nsmb.2149.

17. Guo, H., Ingolia, N.T., Weissman, J.S., and Bartel, D.P. (2010). Mammalian microRNAs predominantly act to decrease target mRNA levels. Nature 466, 835–840. 10.1038/nature09267.

18. Huntzinger, E., Kuzuoglu-Ozturk, D., Braun, J.E., Eulalio, A., Wohlbold, L., and Izaurralde, E. (2013). The interactions of GW182 proteins with PABP and deadenylases are required for both translational repression and degradation of miRNA targets. Nucleic Acids Res 41, 978–994. 10.1093/nar/gks1078.

19. Lazzaretti, D., Tournier, I., and Izaurralde, E. (2009). The C-terminal domains of human TNRC6A, TNRC6B, and TNRC6C silence bound transcripts independently of Argonaute proteins. RNA 15, 1059–1066. 10.1261/rna.1606309.

20. Nishihara, T., Zekri, L., Braun, J.E., and Izaurralde, E. (2013). miRISC recruits decapping factors to miRNA targets to enhance their degradation. Nucleic Acids Res 41, 8692–8705. 10.1093/nar/gkt619.

21. Rehwinkel, J., Behm-Ansmant, I., Gatfield, D., and Izaurralde, E. (2005). A crucial role for GW182 and the DCP1:DCP2 decapping complex in miRNA-mediated gene silencing. RNA 11, 1640–1647. 10.1261/rna.2191905.

22. Till, S., Lejeune, E., Thermann, R., Bortfeld, M., Hothorn, M., Enderle, D., Heinrich, C., Hentze, M.W., and Ladurner, A.G. (2007). A conserved motif in Argonaute-interacting proteins mediates functional interactions through the Argonaute PIWI domain. Nat Struct Mol Biol 14, 897–903. 10.1038/nsmb1302.

23. Mendell, J.T., and Olson, E.N. (2012). MicroRNAs in stress signaling and human disease. Cell 148, 1172–1187. 10.1016/j.cell.2012.02.005.

24. Lujambio, A., and Lowe, S.W. (2012). The microcosmos of cancer. Nature 482, 347–355. 10.1038/nature10888.

25. Calin, G.A., and Croce, C.M. (2006). MicroRNA signatures in human cancers. Nat Rev Cancer 6, 857–866. 10.1038/nrc1997.

26. Gaur, A., Jewell, D.A., Liang, Y., Ridzon, D., Moore, J.H., Chen, C., Ambros, V.R., and Israel, M.A. (2007). Characterization of microRNA expression levels and their biological correlates in human cancer cell lines. Cancer Res 67, 2456–2468. 10.1158/0008-5472.CAN-06-2698.

27. Ventura, A., Young, A.G., Winslow, M.M., Lintault, L., Meissner, A., Erkeland, S.J., Newman, J., Bronson, R.T., Crowley, D., Stone, J.R., et al. (2008). Targeted deletion reveals essential and overlapping functions of the miR-17 through 92 family of miRNA clusters. Cell 132, 875–886. 10.1016/j.cell.2008.02.019.

28. Takamizawa, J., Konishi, H., Yanagisawa, K., Tomida, S., Osada, H., Endoh, H., Harano, T., Yatabe, Y., Nagino, M., Nimura, Y., et al. (2004). Reduced expression of the let-7 microRNAs in human lung cancers in association with shortened postoperative survival. Cancer Res 64, 3753–3756. 10.1158/0008-5472.CAN-04-0637.

29. Calin, G.A., Dumitru, C.D., Shimizu, M., Bichi, R., Zupo, S., Noch, E., Aldler, H., Rattan, S., Keating, M., Rai, K., et al. (2002). Frequent deletions and down-regulation of micro-RNA genes miR15 and miR16 at 13q14 in chronic lymphocytic leukemia. Proc Natl Acad Sci U S A 99, 15524–15529. 10.1073/pnas.242606799.

30. Di Leva, G., and Croce, C.M. (2010). Roles of small RNAs in tumor formation. Trends Mol Med 16, 257–267. 10.1016/j.molmed.2010.04.001.

31. He, L., Thomson, J.M., Hemann, M.T., Hernando-Monge, E., Mu, D., Goodson, S., Powers, S., Cordon-Cardo, C., Lowe, S.W., Hannon, G.J., and Hammond, S.M. (2005). A microRNA polycistron as a potential human oncogene. Nature 435, 828–833. 10.1038/nature03552.

32. Johnson, S.M., Grosshans, H., Shingara, J., Byrom, M., Jarvis, R., Cheng, A., Labourier, E., Reinert, K.L., Brown, D., and Slack, F.J. (2005). RAS is regulated by the let-7 microRNA family. Cell 120, 635–647. 10.1016/j.cell.2005.01.014.

33. Mu, P., Han, Y.C., Betel, D., Yao, E., Squatrito, M., Ogrodowski, P., de Stanchina, E., D’Andrea, A., Sander, C., and Ventura, A. (2009). Genetic dissection of the miR-17∼92 cluster of microRNAs in Myc-induced B-cell lymphomas. Genes Dev 23, 2806–2811. 10.1101/gad.1872909.

34. Merritt, W.M., Lin, Y.G., Han, L.Y., Kamat, A.A., Spannuth, W.A., Schmandt, R., Urbauer, D., Pennacchio, L.A., Cheng, J., Nick, A.M., et al. (2008). Dicer, Drosha, and Outcomes in Patients with Ovarian Cancer. New England Journal of Medicine 359, 2641–2650. DOI 10.1056/NEJMoa0803785.

35. Sand, M., Gambichler, T., Skrygan, M., Sand, D., Scola, N., Altmeyer, P., and Bechara, F.G. (2010). Expression Levels of the microRNA Processing Enzymes Drosha and Dicer in Epithelial Skin Cancer. Cancer Investigation 28, 649–653. 10.3109/07357901003630918.

36. Pampalakis, G., Diamandis, E.P., Katsaros, D., and Sotiropoulou, G. (2010). Down-regulation of dicer expression in ovarian cancer tissues. Clinical Biochemistry 43, 324–327. 10.1016/j.clinbiochem.2009.09.014.

37. Karube, Y., Tanaka, H., Osada, H., Tomida, S., Tatematsu, Y., Yanagisawa, K., Yatabe, Y., Takamizawa, J., Miyoshi, S., Mitsudomi, T., and Takahashi, T. (2005). Reduced expression of Dicer associated with poor prognosis in lung cancer patients. Cancer Science 96, 111–115. DOI 10.1111/j.1349-7006.2005.00015.x.

38. Lin, R.J., Lin, Y.C., Chen, J., Kuo, H.H., Chen, Y.Y., Diccianni, M.B., London, W.B., Chang, C.H., and Yu, A.L. (2010). microRNA Signature and Expression of and Can Predict Prognosis and Delineate Risk Groups in Neuroblastoma. Cancer Res 70, 7841–7850. 10.1158/0008-5472.Can-10-0970.

39. Kumar, M.S., Lu, J., Mercer, K.L., Golub, T.R., and Jacks, T. (2007). Impaired microRNA processing enhances cellular transformation and tumorigenesis. Nat Genet 39, 673–677. 10.1038/ng2003.

40. Anglesio, M.S., Wang, Y., Yang, W., Senz, J., Wan, A., Heravi-Moussavi, A., Salamanca, C., Maines-Bandiera, S., Huntsman, D.G., and Morin, G.B. (2013). Cancer-associated somatic DICER1 hotspot mutations cause defective miRNA processing and reverse-strand expression bias to predominantly mature 3p strands through loss of 5p strand cleavage. J Pathol 229, 400–409. 10.1002/path.4135.

41. Heravi-Moussavi, A., Anglesio, M.S., Cheng, S.W., Senz, J., Yang, W., Prentice, L., Fejes, A.P., Chow, C., Tone, A., Kalloger, S.E., et al. (2012). Recurrent somatic DICER1 mutations in nonepithelial ovarian cancers. N Engl J Med 366, 234–242. 10.1056/NEJMoa1102903.

42. Rakheja, D., Chen, K.S., Liu, Y.J., Shukla, A.A., Schmid, V., Chang, T.C., Khokhar, S., Wickiser, J.E., Karandikar, N.J., Malter, J.S., et al. (2017). Somatic mutations in DROSHA and DICER1 impair microRNA biogenesis through distinct mechanisms in Wilms tumours. Nature Communications 8. ARTN 1617710.1038/ncomms16177.

43. Melo, S.A., Moutinho, C., Ropero, S., Calin, G.A., Rossi, S., Spizzo, R., Fernandez, A.F., Davalos, V., Villanueva, A., Montoya, G., et al. (2010). A Genetic Defect in Exportin-5 Traps Precursor MicroRNAs in the Nucleus of Cancer Cells. Cancer Cell 18, 303–315. 10.1016/j.ccr.2010.09.007.

44. Bronisz, A., Rooj, A.K., Krawczynski, K., Peruzzi, P., Salinska, E., Nakano, I., Purow, B., Chiocca, E.A., and Godlewski, J. (2020). The nuclear DICER-circular RNA complex drives the deregulation of the glioblastoma cell microRNAome. Sci Adv 6. ARTN eabc022110.1126/sciadv.abc0221.

45. Martello, G., Rosato, A., Ferrari, F., Manfrin, A., Cordenonsi, M., Dupont, S., Enzo, E., Guzzardo, V., Rondina, M., Spruce, T., et al. (2010). A MicroRNA targeting dicer for metastasis control. Cell 141, 1195–1207. 10.1016/j.cell.2010.05.017.

46. Ramirez-Moya, J., Wert-Lamas, L., Riesco-Eizaguirre, G., and Santisteban, P. (2019). Impaired microRNA processing by DICER1 downregulation endows thyroid cancer with increased aggressiveness. Oncogene 38, 5486–5499. 10.1038/s41388-019-0804-8.

47. Chang, T.C., Yu, D.N., Lee, Y.S., Wentzel, E.A., Arking, D.E., West, K.M., Dang, C.V., Thomas-Tikhonenko, A., and Mendell, J.T. (2008). Widespread microRNA repression by Myc contributes to tumorigenesis. Nat Genet 40, 43–50. 10.1038/ng.2007.30.

48. Godlewski, J., Ferrer-Luna, R., Rooj, A.K., Mineo, M., Ricklefs, F., Takeda, Y.S., Nowicki, M.O., Salinska, E., Nakano, I., Lee, H., et al. (2017). MicroRNA Signatures and Molecular Subtypes of Glioblastoma: The Role of Extracellular Transfer. Stem Cell Reports 8, 1497–1505. 10.1016/j.stemcr.2017.04.024.

49. Lu, J., Getz, G., Miska, E.A., Alvarez-Saavedra, E., Lamb, J., Peck, D., Sweet-Cordero, A., Ebert, B.L., Mak, R.H., Ferrando, A.A., et al. (2005). MicroRNA expression profiles classify human cancers. Nature 435, 834–838. 10.1038/nature03702.

50. Lujambic, A., Ropero, S., Ballestar, E., Fraga, M.F., Cerrato, C., Setién, F., Casado, S., Suarez-Gauthier, A., Sanchez-Cespedes, M., Git, A., et al. (2007). Genetic unmasking of an epigenetically silenced microRNA in human cancer cells (vol 67, pg 1424, 2007). Cancer Res 67, 3492–3492.

51. Thomson, J.M., Newman, M., Parker, J.S., Morin-Kensicki, E.M., Wright, T., and Hammond, S.M. (2006). Extensive post-transcriptional regulation of microRNAs and its implications for cancer. Genes Dev 20, 2202–2207. 10.1101/gad.1444406.

52. Ozen, M., Creighton, C.J., Ozdemir, M., and Ittmann, M. (2008). Widespread deregulation of microRNA expression in human prostate cancer. Oncogene 27, 1788–1793. 10.1038/sj.onc.1210809.

53. Chiosea, S., Jelezcova, E., Chandran, U., Acquafondata, M., McHale, T., Sobol, R.W., and Dhir, R. (2006). Up-regulation of dicer, a component of the microRNA machinery, in prostate adenocarcinoma. American Journal of Pathology 169, 1812–1820. 10.2353/ajpath.2006.060480.

54. Jakymiw, A., Patel, R.S., Deming, N., Bhattacharyya, I., Shah, P., Lamont, R.J., Stewart, C.M., Cohen, D.M., and Chan, E.K.L. (2010). Overexpression of Dicer as a Result of Reduced let-7 MicroRNA Levels Contributes to Increased Cell Proliferation of Oral Cancer Cells. Genes Chromosomes & Cancer 49, 549–559. 10.1002/gcc.20765.

55. Martin, M.G., Payton, J.E., and Link, D.C. (2009). Dicer and outcomes in patients with acute myeloid leukemia (AML). Leuk Res 33, e127. 10.1016/j.leukres.2009.02.003.

56. Muralidhar, B., Goldstein, L.D., Ng, G., Winder, D.M., Palmer, R.D., Gooding, E.L., Barbosa-Morais, N.L., Mukherjee, G., Thorne, N.P., Roberts, I., et al. (2007). Global microRNA profiles in cervical squamous cell carcinoma depend on Drosha expression levels. Journal of Pathology 212, 368–377. 10.1002/path.2179.

57. Sugito, N., Ishiguro, H., Kuwabara, Y., Kimura, M., Mitsui, A., Kurehara, H., Ando, T., Mori, R., Takashima, N., Ogawa, R., and Fujii, Y. (2006). RNASEN regulates cell proliferation and affects survival in esophageal cancer patients. Clinical Cancer Research 12, 7322–7328. 10.1158/1078-0432.Ccr-06-0515.

58. Faber, C., Horst, D., Hlubek, F., and Kirchner, T. (2011). Overexpression of Dicer predicts poor survival in colorectal cancer. Eur J Cancer 47, 1414–1419. 10.1016/j.ejca.2011.01.006.

59. Kumar, M.S., Pester, R.E., Chen, C.Y., Lane, K., Chin, C., Lu, J., Kirsch, D.G., Golub, T.R., and Jacks, T. (2009). Dicer1 functions as a haploinsufficient tumor suppressor. Genes & Development 23, 2700–2704. 10.1101/gad.1848209.

60. Lambertz, I., Nittner, D., Mestdagh, P., Denecker, G., Vandesompele, J., Dyer, M.A., and Marine, J.C. (2010). Monoallelic but not biallelic loss of Dicer1 promotes tumorigenesis in vivo. Cell Death Differ 17, 633–641. 10.1038/cdd.2009.202.

61. Mito, J.K., Min, H.D., Ma, Y., Carter, J.E., Brigman, B.E., Dodd, L., Dankort, D., McMahon, M., and Kirsch, D.G. (2013). Oncogene-dependent control of miRNA biogenesis and metastatic progression in a model of undifferentiated pleomorphic sarcoma. J Pathol 229, 132–140. 10.1002/path.4099.

62. Wegert, J., Ishaque, N., Vardapour, R., Georg, C., Gu, Z., Bieg, M., Ziegler, B., Bausenwein, S., Nourkami, N., Ludwig, N., et al. (2015). Mutations in the SIX1/2 pathway and the DROSHA/DGCR8 miRNA microprocessor complex underlie high-risk blastemal type Wilms tumors. Cancer Cell 27, 298–311. 10.1016/j.ccell.2015.01.002.

63. Walz, A.L., Ooms, A., Gadd, S., Gerhard, D.S., Smith, M.A., Guidry Auvil, J.M., Meerzaman, D., Chen, Q.R., Hsu, C.H., Yan, C., et al. (2015). Recurrent DGCR8, DROSHA, and SIX homeodomain mutations in favorable histology Wilms tumors. Cancer Cell 27, 286–297. 10.1016/j.ccell.2015.01.003.

64. Johnson, K.C., Johnson, S.T., Liu, J., Chu, Y., Arana, C., Han, Y., Wang, T., and Corey, D.R. (2023). Consequences of depleting TNRC6, AGO, and DROSHA proteins on expression of microRNAs. RNA 29, 1166–1184. 10.1261/rna.079647.123.

65. Kim, Y.K., Kim, B., and Kim, V.N. (2016). Re-evaluation of the roles of DROSHA, Export in 5, and DICER in microRNA biogenesis. Proc Natl Acad Sci U S A 113, E1881–1889. 10.1073/pnas.1602532113.

66. Cheloufi, S., Dos Santos, C.O., Chong, M.M., and Hannon, G.J. (2010). A dicer-independent miRNA biogenesis pathway that requires Ago catalysis. Nature 465, 584–589. 10.1038/nature09092.

67. Chong, M.M., Zhang, G., Cheloufi, S., Neubert, T.A., Hannon, G.J., and Littman, D.R. (2010). Canonical and alternate functions of the microRNA biogenesis machinery. Genes Dev 24, 1951–1960. 10.1101/gad.1953310.

68. Cifuentes, D., Xue, H., Taylor, D.W., Patnode, H., Mishima, Y., Cheloufi, S., Ma, E., Mane, S., Hannon, G.J., Lawson, N.D., et al. (2010). A novel miRNA processing pathway independent of Dicer requires Argonaute2 catalytic activity. Science 328, 1694–1698. 10.1126/science.1190809.

69. Okamura, K., Hagen, J.W., Duan, H., Tyler, D.M., and Lai, E.C. (2007). The mirtron pathway generates microRNA-class regulatory RNAs in Drosophila. Cell 130, 89–100. 10.1016/j.cell.2007.06.028.

70. Ruby, J.G., Jan, C.H., and Bartel, D.P. (2007). Intronic microRNA precursors that bypass Drosha processing. Nature 448, 83–86. 10.1038/nature05983.

71. Yang, J.S., and Lai, E.C. (2011). Alternative miRNA biogenesis pathways and the interpretation of core miRNA pathway mutants. Mol Cell 43, 892–903. 10.1016/j.molcel.2011.07.024.

72. Vechetti, I.J., Jr., Wen, Y., Chaillou, T., Murach, K.A., Alimov, A.P., Figueiredo, V.C., Dal-Pai-Silva, M., and McCarthy, J.J. (2019). Life-long reduction in myomiR expression does not adversely affect skeletal muscle morphology. Sci Rep 9, 5483. 10.1038/s41598-019-41476-8.

73. Olejniczak, S.H., La Rocca, G., Radler, M.R., Egan, S.M., Xiang, Q., Garippa, R., and Thompson, C.B. (2016). Coordinated Regulation of Cap-Dependent Translation and MicroRNA Function by Convergent Signaling Pathways. Mol Cell Biol 36, 2360–2373. 10.1128/MCB.01011-15.

74. Golden, R.J., Chen, B., Li, T., Braun, J., Manjunath, H., Chen, X., Wu, J., Schmid, V., Chang, T.C., Kopp, F., et al. (2017). An Argonaute phosphorylation cycle promotes microRNA-mediated silencing. Nature 542, 197–202. 10.1038/nature21025.

75. La Rocca, G., Olejniczak, S.H., Gonzalez, A.J., Briskin, D., Vidigal, J.A., Spraggon, L., DeMatteo, R.G., Radler, M.R., Lindsten, T., Ventura, A., et al. (2015). In vivo, Argonaute-bound microRNAs exist predominantly in a reservoir of low molecular weight complexes not associated with mRNA. Proc Natl Acad Sci U S A 112, 767–772. 10.1073/pnas.1424217112.

76. Quevillon Huberdeau, M., Zeitler, D.M., Hauptmann, J., Bruckmann, A., Fressigne, L., Danner, J., Piquet, S., Strieder, N., Engelmann, J.C., Jannot, G., et al. (2017). Phosphorylation of Argonaute proteins affects mRNA binding and is essential for microRNA-guided gene silencing in vivo. EMBO J 36, 2088–2106. 10.15252/embj.201696386.

77. Shah, V.N., Neumeier, J., Huberdeau, M.Q., Zeitler, D.M., Bruckmann, A., Meister, G., and Simard, M.J. (2023). Casein kinase 1 and 2 phosphorylate Argonaute proteins to regulate miRNA-mediated gene silencing. EMBO Rep 24, e57250. 10.15252/embr.202357250.

78. Shui, B., Beyett, T.S., Chen, Z., Li, X., La Rocca, G., Gazlay, W.M., Eck, M.J., Lau, K.S., Ventura, A., and Haigis, K.M. (2023). Oncogenic K-Ras suppresses global miRNA function. Mol Cell 83, 2509–2523 e2513. 10.1016/j.molcel.2023.06.008.

79. Fukagawa, T., Nogami, M., Yoshikawa, M., Ikeno, M., Okazaki, T., Takami, Y., Nakayama, T., and Oshimura, M. (2004). Dicer is essential for formation of the heterochromatin structure in vertebrate cells. Nat Cell Biol 6, 784–791. 10.1038/ncb1155.

80. Giles, K.E., Ghirlando, R., and Felsenfeld, G. (2010). Maintenance of a constitutive heterochromatin domain in vertebrates by a Dicer-dependent mechanism. Nat Cell Biol 12, 94–99; sup pp 91–96. 10.1038/ncb2010.

81. Gullerova, M., and Proudfoot, N.J. (2012). Convergent transcription induces transcriptional gene silencing in fission yeast and mammalian cells. Nat Struct Mol Biol 19, 1193–1201. 10.1038/nsmb.2392.

82. Okamura, K., and Lai, E.C. (2008). Endogenous small interfering RNAs in animals. Nat Rev Mol Cell Biol 9, 673–678. 10.1038/nrm2479.

83. Song, M.S., and Rossi, J.J. (2017). Molecular mechanisms of Dicer: endonuclease and enzymatic activity. Biochem J 474, 1603–1618. 10.1042/BCJ20160759.

84. Tam, O.H., Aravin, A.A., Stein, P., Girard, A., Murchison, E.P., Cheloufi, S., Hodges, E., Anger, M., Sachidanandam, R., Schultz, R.M., and Hannon, G.J. (2008). Pseudogene-derived small interfering RNAs regulate gene expression in mouse oocytes. Nature 453, 534–538. 10.1038/nature06904.

85. Kaneko, H., Dridi, S., Tarallo, V., Gelfand, B.D., Fowler, B.J., Cho, W.G., Kleinman, M.E., Ponicsan, S.L., Hauswirth, W.W., Chiodo, V.A., et al. (2011). DICER1 deficit induces Alu RNA toxicity in age-related macular degeneration. Nature 471, 325–330. 10.1038/nature09830.

86. Francia, S., Michelini, F., Saxena, A., Tang, D., de Hoon, M., Anelli, V., Mione, M., Carninci, P., and d’Adda di Fagagna, F. (2012). Site-specific DICER and DROSHA RNA products control the DNA-damage response. Nature 488, 231–235. 10.1038/nature11179.

87. Michelini, F., Pitchiaya, S., Vitelli, V., Sharma, S., Gioia, U., Pessina, F., Cabrini, M., Wang, Y., Capozzo, I., Iannelli, F., et al. (2017). Damage-induced lncRNAs control the DNA damage response through interaction with DDRNAs at individual double-strand breaks. Nat Cell Biol 19, 1400–1411. 10.1038/ncb3643.

88. Cirera-Salinas, D., Yu, J., Bodak, M., Ngondo, R.P., Herbert, K.M., and Ciaudo, C. (2017). Noncanonical function of DGCR8 controls mESC exit from pluripotency. J Cell Biol 216, 355–366. 10.1083/jcb.201606073.

89. Macias, S., Cordiner, R.A., Gautier, P., Plass, M., and Caceres, J.F. (2015). DGCR8 Acts as an Adaptor for the Exosome Complex to Degrade Double-Stranded Structured RNAs. Mol Cell 60, 873–885. 10.1016/j.molcel.2015.11.011.

90. Sala, L., Kumar, M., Prajapat, M., Chandrasekhar, S., Cosby, R.L., La Rocca, G., Macfarlan, T.S., Awasthi, P., Chari, R., Kruhlak, M., and Vidigal, J.A. (2023). AGO2 silences mobile transposons in the nucleus of quiescent cells. Nat Struct Mol Biol 30, 1985–1995. 10.1038/s41594-023-01151-z.

91. La Rocca, G., and Cavalieri, V. (2022). Roles of the Core Components of the Mammalian miRISC in Chromatin Biology. Genes (Basel) 13. 10.3390/genes13030414.

92. Lu, W.T., Hawley, B.R., Skalka, G.L., Baldock, R.A., Smith, E.M., Bader, A.S., Malewicz, M., Watts, F.Z., Wilczynska, A., and Bushell, M. (2018). Drosha drives the formation of DNA:RNA hybrids around DNA break sites to facilitate DNA repair. Nat Commun 9, 532. 10.1038/s41467-018-02893-x.

93. Castel, S.E., Ren, J., Bhattacharjee, S., Chang, A.Y., Sanchez, M., Valbuena, A., Antequera, F., and Martienssen, R.A. (2014). Dicer promotes transcription termination at sites of replication stress to maintain genome stability. Cell 159, 572–583. 10.1016/j.cell.2014.09.031.

94. La Rocca, G., King, B., Shui, B., Li, X.Y., Zhang, M.S., Akat, K.M., Ogrodowski, P., Mastroleo, C., Chen, K., Cavalieri, V., et al. (2021). Inducible and reversible inhibition of miRNA-mediated gene repression in vivo. Elife 10. ARTN e7094810.7554/eLife.70948

95. Han, Y.C., Vidigal, J.A., Mu, P., Yao, E., Singh, I., Gonzalez, A.J., Concepcion, C.P., Bonetti, C., Ogrodowski, P., Carver, B., et al. (2015). An allelic series of miR-17 approximately 92-mutant mice uncovers functional specialization and cooperation among members of a microRNA polycistron. Nat Genet 47, 766–775. 10.1038/ng.3321.

96. Yang, Z., Jakymiw, A., Wood, M.R., Eystathioy, T., Rubin, R.L., Fritzler, M.J., and Chan, E.K. (2004). GW182 is critical for the stability of GW bodies expressed during the cell cycle and cell proliferation. J Cell Sci 117, 5567–5578. 10.1242/jcs.01477.

97. Hanahan, D., and Weinberg, R.A. (2011). Hallmarks of cancer: the next generation. Cell 144, 646–674. 10.1016/j.cell.2011.02.013.

98. Adams, J.M., Harris, A.W., Pinkert, C.A., Corcoran, L.M., Alexander, W.S., Cory, S., Palmiter, R.D., and Brinster, R.L. (1985). The c-myc oncogene driven by immunoglobulin enhancers induces lymphoid malignancy in transgenic mice. Nature 318, 533–538. 10.1038/318533a0.

99. Maddalo, D., Manchado, E., Concepcion, C.P., Bonetti, C., Vidigal, J.A., Han, Y.C., Ogrodowski, P., Crippa, A., Rekhtman, N., de Stanchina, E., et al. (2014). In vivo engineering of oncogenic chromosomal rearrangements with the CRISPR/Cas9 system. Nature 516, 423–427. 10.1038/nature13902.

100. DuPage, M., Dooley, A.L., and Jacks, T. (2009). Conditional mouse lung cancer models using adenoviral or lentiviral delivery of Cre recombinase. Nat Protoc 4, 1064–1072. 10.1038/nprot.2009.95.

101. Kirsch, D.G., Dinulescu, D.M., Miller, J.B., Grimm, J., Santiago, P.M., Young, N.P., Nielsen, G.P., Quade, B.J., Chaber, C.J., Schultz, C.P., et al. (2007). A spatially and temporally restricted mouse model of soft tissue sarcoma. Nat Med 13, 992–997. 10.1038/nm1602.

102. Sekar, V., Marmol-Sanchez, E., Kalogeropoulos, P., Stanicek, L., Sagredo, E.A., Widmark, A., Doukoumopoulos, E., Bonath, F., Biryukova, I., and Friedlander, M.R. (2023). Detection of transcriptome-wide microRNA-target interactions in single cells with agoTRIBE. Nat Biotechnol. 10.1038/s41587-023-01951-0.

103. Hauptmann, J., Schraivogel, D., Bruckmann, A., Manickavel, S., Jakob, L., Eichner, N., Pfaff, J., Urban, M., Sprunck, S., Hafner, M., et al. (2015). Biochemical isolation of Argonaute protein complexes by Ago-APP. Proc Natl Acad Sci U S A 112, 11841–11845. 10.1073/pnas.1506116112.

104. Briskin, D., Wang, P.Y., and Bartel, D.P. (2020). The biochemical basis for the cooperative action of microRNAs. Proc Natl Acad Sci U S A 117, 17764–17774. 10.1073/pnas.1920404117.

105. Welte, T., Goulois, A., Stadler, M.B., Hess, D., Soneson, C., Neagu, A., Azzi, C., Wisser, M.J., Seebacher, J., Schmidt, I., et al. (2023). Convergence of multiple RNA-silencing pathways on GW182/TNRC6. Mol Cell 83, 2478–2492 e2478. 10.1016/j.molcel.2023.06.001.

106. Danner, J., Pai, B., Wankerl, L., and Meister, G. (2017). Peptide-Based Inhibition of miRNA-Guided Gene Silencing. Drug Target Mirna: Methods and Protocols 1517, 199–210. 10.1007/978-1-4939-6563-2_14.

107. Li, X., Pritykin, Y., Concepcion, C.P., Lu, Y., La Rocca, G., Zhang, M., King, B., Cook, P.J., Au, Y.W., Popow, O., et al. (2020). High-Resolution In Vivo Identification of miRNA Targets by Halo-Enhanced Ago2 Pull-Down. Mol Cell 79, 167–179 e111. 10.1016/j.molcel.2020.05.009.

108. Li, Y., Wang, L., Rivera-Serrano, E.E., Chen, X., and Lemon, S.M. (2019). TNRC6 proteins modulate hepatitis C virus replication by spatially regulating the binding of miR-122/Ago2 complexes to viral RNA. Nucleic Acids Res 47, 6411–6424. 10.1093/nar/gkz278.

109. Zolboot, N., Xiao, Y., Du, J.X., Ghanem, M.M., Choi, S.Y., Junn, M.J., Zampa, F., Huang, Z., MacRae, I.J., and Lippi, G. (2023). MicroRNAs are necessary for the emergence of Purkinje cell identity. bioRxiv. 10.1101/2023.09.28.560023.

110. Molina-Sanchez, P., Ruiz de Galarreta, M., Yao, M.A., Lindblad, K.E., Bresnahan, E., Bitterman, E., Martin, T.C., Rubenstein, T., Nie, K., Golas, J., et al. (2020). Cooperation Between Distinct Cancer Driver Genes Underlies Intertumor Heterogeneity in Hepatocellular Carcinoma. Gastroenterology 159, 2203–2220 e2214. 10.1053/j.gastro.2020.08.015.

111. Ivics, Z., Hackett, P.B., Plasterk, R.H., and Izsvak, Z. (1997). Molecular reconstruction of Sleeping Beauty, a Tc1-like transposon from fish, and its transposition in human cells. Cell 91, 501–510. 10.1016/s0092-8674(00)80436-5.

112. Licursi, V., Conte, F., Fiscon, G., and Paci, P. (2019). MIENTURNET: an interactive web tool for microRNA-target enrichment and network-based analysis. BMC Bioinformatics 20, 545. 10.1186/s12859-019-3105-x.

113. Lessel, D., Zeitler, D.M., Reijnders, M.R.F., Kazantsev, A., Hassani Nia, F., Bartholomaus, A., Martens, V., Bruckmann, A., Graus, V., McConkie-Rosell, A., et al. (2020). Germline AGO2 mutations impair RNA interference and human neurological development. Nat Commun 11, 5797. 10.1038/s41467-020-19572-5.

114. Antoccia, A., Degrassi, F., Battistoni, A., Ciliutti, P., and Tanzarella, C. (1991). In vitro micronucleus test with kinetochore staining: evaluation of test performance. Mutagenesis 6, 319–324. 10.1093/mutage/6.4.319.

115. Sundararajan, K., and Straight, A.F. (2022). Centromere Identity and the Regulation of Chromosome Segregation. Front Cell Dev Biol 10, 914249. 10.3389/fcell.2022.914249.

116. Prosser, S.L., and Pelletier, L. (2017). Mitotic spindle assembly in animal cells: a fine balancing act. Nat Rev Mol Cell Biol 18, 187–201. 10.1038/nrm.2016.162.

117. Lens, S.M.A., and Medema, R.H. (2019). Cytokinesis defects and cancer. Nat Rev Cancer 19, 32–45. 10.1038/s41568-018-0084-6.

118. Janssen, A., van der Burg, M., Szuhai, K., Kops, G.J., and Medema, R.H. (2011). Chromosome segregation errors as a cause of DNA damage and structural chromosome aberrations. Science 333, 1895–1898. 10.1126/science.1210214.

119. Li, R., and Zhu, J. (2022). Effects of aneuploidy on cell behaviour and function. Nat Rev Mol Cell Biol 23, 250–265. 10.1038/s41580-021-00436-9.

120. Sparr, C., and Meitinger, F. (2025). Prolonged mitosis: A key indicator for detecting stressed and damaged cells. Curr Opin Cell Biol 92, 102449. 10.1016/j.ceb.2024.102449.

121. Vitale, I., Galluzzi, L., Castedo, M., and Kroemer, G. (2011). Mitotic catastrophe: a mechanism for avoiding genomic instability. Nat Rev Mol Cell Biol 12, 385–392. 10.1038/nrm3115.

122. Hochegger, H., Takeda, S., and Hunt, T. (2008). Cyclin-dependent kinases and cell-cycle transitions: does one fit all? Nat Rev Mol Cell Biol 9, 910–916. 10.1038/nrm2510.

123. Chang, D.C., Xu, N., and Luo, K.Q. (2003). Degradation of cyclin B is required for the onset of anaphase in Mammalian cells. J Biol Chem 278, 37865–37873. 10.1074/jbc.M306376200.

124. Morgan, M.A., and Lawrence, T.S. (2015). Molecular Pathways: Overcoming Radiation Resistance by Targeting DNA Damage Response Pathways. Clin Cancer Res 21, 2898–2904. 10.1158/1078-0432.CCR-13-3229.

125. Zhang, M., Qiu, Q., Li, Z., Sachdeva, M., Min, H., Cardona, D.M., DeLaney, T.F., Han, T., Ma, Y., Luo, L., et al. (2015). HIF-1 Alpha Regulates the Response of Primary Sarcomas to Radiation Therapy through a Cell Autonomous Mechanism. Radiat Res 183, 594–609. 10.1667/RR14016.1.

126. Wisdom, A.J., Hong, C.S., Lin, A.J., Xiang, Y., Cooper, D.E., Zhang, J., Xu, E.S., Kuo, H.C., Mowery, Y.M., Carpenter, D.J., et al. (2019). Neutrophils promote tumor resistance to radiation therapy. Proc Natl Acad Sci U S A 116, 18584–18589. 10.1073/pnas.1901562116.

127. Floyd, W., Pierpoint, M., Su, C., Patel, R., Luo, L., Deland, K., Wisdom, A.J., Zhu, D., Ma, Y., DeWitt, S.B., et al. (2023). Atrx deletion impairs CGAS/STING signaling and increases sarcoma response to radiation and oncolytic herpesvirus. J Clin Invest 133. 10.1172/JCI149310.

128. Liu, H., Arsie, R., Schwabe, D., Schilling, M., Minia, I., Alles, J., Boltengagen, A., Kocks, C., Falcke, M., Friedman, N., et al. (2023). SLAM-Drop-seq reveals mRNA kinetic rates throughout the cell cycle. Mol Syst Biol 19, 1–23. 10.15252/msb.202211427.

129. Shen, M., and Kang, Y. (2023). Cancer fitness genes: emerging therapeutic targets for metastasis. Trends Cancer 9, 69–82. 10.1016/j.trecan.2022.08.007.

130. Wang, G., Huang, Y., Wu, Z., Zhao, C., Cong, H., Ju, S., and Wang, X. (2019). KRAS-mutant colon cancer cells respond to combined treatment of ABT263 and axitinib. Biosci Rep 39. 10.1042/BSR20181786.

131. Nelson, P.T., De Planell-Saguer, M., Lamprinaki, S., Kiriakidou, M., Zhang, P., O’Doherty, U., and Mourelatos, Z. (2007). A novel monoclonal antibody against human Argonaute proteins reveals unexpected characteristics of miRNAs in human blood cells. RNA 13, 1787–1792. 10.1261/rna.646007.

132. Dobin, A., Davis, C.A., Schlesinger, F., Drenkow, J., Zaleski, C., Jha, S., Batut, P., Chaisson, M., and Gingeras, T.R. (2013). STAR: ultrafast universal RNA-seq aligner. Bioinformatics 29, 15–21. 10.1093/bioinformatics/bts635.

133. Love, M.I., Huber, W., and Anders, S. (2014). Moderated estimation of fold change and dispersion for RNA-seq data with DESeq2. Genome Biol 15, 550. 10.1186/s13059-014-0550-8.

